# Compartment-dependent chromatin interaction dynamics revealed by liquid chromatin Hi-C

**DOI:** 10.1101/704957

**Authors:** Houda Belaghzal, Tyler Borrman, Andrew D. Stephens, Denis L. Lafontaine, Sergey V. Venev, Zhiping Weng, John F. Marko, Job Dekker

## Abstract

Chromosomes are folded so that active and inactive chromatin domains are spatially segregated. Compartmentalization is thought to occur through polymer phase/microphase separation mediated by interactions between loci of similar type. The nature and dynamics of these interactions are not known. We developed liquid chromatin Hi-C to map the stability of associations between loci. Before fixation and Hi-C, chromosomes are fragmented removing the strong polymeric constraint to enable detection of intrinsic locus-locus interaction stabilities. Compartmentalization is stable when fragments are over 10-25 kb. Fragmenting chromatin into pieces smaller than 6 kb leads to gradual loss of genome organization. Dissolution kinetics of chromatin interactions vary for different chromatin domains. Lamin-associated domains are most stable, while interactions among speckle and polycomb-associated loci are more dynamic. Cohesin-mediated loops dissolve after fragmentation, possibly because cohesin rings slide off nearby DNA ends. Liquid chromatin Hi-C provides a genome-wide view of chromosome interaction dynamics.

**Highlights:**

- Liquid chromatin Hi-C detects chromatin interaction dissociation rates genome-wide
- Chromatin conformations in distinct nuclear compartments differ in stability
- Stable heterochromatic associations are major drivers of chromatin phase separation
- CTCF-CTCF loops are stabilized by encirclement of loop bases by cohesin rings

## INTRODUCTION

The spatial organization of chromosomes plays roles in many aspects of genome function, including gene regulation, DNA replication, DNA repair, and chromosome compaction and segregation. Genomic and imaging approaches are producing high-resolution descriptions of the conformation of chromosomes in cell populations, in single cells, across the cell cycle, and during development (Bickmore and van Steensel, 2013; Bonev and Cavalli, 2016; Chen et al., 2018; Dekker et al., 2017; Dekker and Mirny, 2016; Gibcus et al., 2018; Hug et al., 2017; Kaaij et al., 2018; Lieberman-Aiden et al., 2009; Nagano et al., 2013; Nagano et al., 2017; Naumova et al., 2013; Nir et al., 2018; Ramani et al., 2017; Rao et al., 2014; Wang et al., 2016).

At the scale of the nucleus, chromosomes occupy separate territories with some interdigitation where they touch (Branco and Pombo, 2006; Cremer and Cremer, 2001). At the mega-base (Mb) scale, different types of chromosomal domains can be discerned. Domains of silent heterochromatin are peripherally localized near the nuclear lamina, and around the nucleolus (Bickmore and van Steensel, 2013; Guelen et al., 2008; Padeken and Heun, 2014). Other domains of silent chromatin such as those bound by polycomb complexes can cluster together as well, e.g. to form polycomb bodies (Pirrotta and Li, 2012). Active segments of the genome are found around bodies such as nuclear speckles (Fakan and Puvion, 1980; Spector and Lamond, 2011) and structures enriched in transcription regulatory factors (Cho et al., 2018; Chong et al., 2018; Iborra et al., 1996; Pombo et al., 1999).

Hi-C data confirmed the spatial segregation of active and inactive segments of chromosomes: chromatin interaction maps display a “plaid” pattern, which reflects the segregation of the genome in two major spatial compartments referred to as A and B compartments that correspond to open and active chromatin and closed and silent chromatin, respectively (Lieberman-Aiden et al., 2009; Simonis et al., 2006). High-resolution (kb) Hi-C maps allowed splitting these two major compartments in 5 subtypes (A1, A2, B1, B2, and B3) that differ in interaction patterns and chromatin modification status (Rao et al., 2014). In mammalian cells these domains are typically hundreds of kb up to several Mb in size. Belmont and colleagues related these sub-compartments to structures such as speckles (A1 domains) in the interior and the peripheral lamina (B2 and B3) (Chen et al., 2018). Data obtained with SPRITE showed similar clustering of genomic sub-compartments around the nucleolus, speckles, and the periphery (Quinodoz et al., 2018).

At the scale of hundreds of kb, Hi-C and 5C experiments uncovered topologically associating domains (TADs; (Dixon et al., 2012; Nora et al., 2012)). TADs were identified as self-interacting domains separated by boundaries that are in many cases bound by CTCF. Higher resolution Hi-C (Rao et al., 2014), ChIA-PET (Tang et al., 2015), and 4C data (de Wit et al., 2015; Guo et al., 2015; Vietri Rudan et al., 2015) showed that CTCF sites at boundaries can engage in looping interactions, especially when the sites are in convergent orientation.

Major new questions revolve around the molecular and biophysical processes by which different aspects of chromosome conformation form. Significant progress has been made in developing and testing mechanistic models for TAD and loop formation. The model that currently has most experimental support proposes that TADs and loops form via loop extrusion performed by the cohesin complex (Alipour and Marko, 2012; Fudenberg et al., 2017; Fudenberg et al., 2016; Nasmyth, 2001; Riggs, 1990; Sanborn et al., 2015). In this model, loop extrusion proceeds till the complex encounters a CTCF protein bound to a CTCF site in the convergent orientation. As a result, CTCF-CTCF loops form, and interactions between loci located between convergent CTCF sites are generally increased (TADs). The mechanisms of loop extrusion are not known in detail yet, but may involve the cohesin ring encircling strands of DNA at the bases of loops in a topological or pseudo-topological manner (Fudenberg et al., 2017; Haering et al., 2008; Ivanov and Nasmyth, 2005; Kagey et al., 2010; Srinivasan et al., 2018).

Much less is known about the processes that determine nuclear compartmentalization. Mechanisms of compartmentalization are distinct from the formation of TADs and loops, as mutations in CTCF or cohesin disrupt TADs but not compartmentalization per se (Haarhuis et al., 2017; Nora et al., 2017; Nuebler et al., 2018; Rao et al., 2017; Schwarzer et al., 2017; Wutz et al., 2017). Compartmentalization has been proposed to be the result of polymer phase separation driven by attractions between chromatin domains of the same or similar status (Di Pierro et al., 2016; Erdel and Rippe, 2018; Falk et al., 2019; Jost et al., 2014; Lieberman-Aiden et al., 2009; MacPherson et al., 2018; Michieletto et al., 2016; Nuebler et al., 2018; Shi et al., 2018). Polymer models simulating such attractions can reproduce the plaid pattern characteristic of Hi-C interaction maps (Di Pierro et al., 2016; Falk et al., 2019; Jost et al., 2014; Nuebler et al., 2018). However, the molecular and biophysical basis of these attractions is unknown. Possibly these attractions result from co-association of domains with sub-nuclear bodies that themselves appear to form by a process of liquid-liquid phase separation (e.g. (Courchaine et al., 2016; Feric et al., 2016; Larson et al., 2017; Marzahn et al., 2016; Sawyer et al., 2018; Strom et al., 2017; Zhu and Brangwynne, 2015). An example is the interaction between heterochromatic loci driven by multivalent interactions among HP1 proteins and between HP1 proteins and H3K9me3-modified chromatin domains (Larson et al., 2017; Strom et al., 2017).

Hi-C contact maps readily reveal global spatial separation of active and inactive chromatin domains, and Hi-C sub-compartments suggest the presence of a number of different types of sub-nuclear neighborhoods. However, Hi-C interaction maps are population-averaged, steady-state datasets and do not reveal the biophysical nature of the interactions that lead to this diverse set of nuclear neighborhoods or the dynamic mobility of loci within these compartments. For instance, A and B compartments appear equally prominent in Hi-C datasets, but whether the forces that lead to their formation are equally strong, frequent, or dynamic is not known.

Live-cell imaging studies have shown that loci are constrained in their motion and that there are substantial variations in the dynamics and mobility of different loci, e.g. euchromatic vs. heterochromatic loci and loci tethered to the nuclear periphery vs. loci located in the nuclear interior (Bronshtein et al., 2009; Bronshtein et al., 2015; Hediger et al., 2002; Marshall et al., 1997; Nagashima et al., 2019; Shinkai et al., 2016; Thakar et al., 2006; Therizols et al., 2010). Such differences can be reproduced in coarse-grained simulations of chromatin (Liu et al., 2018). Imaging-based studies have been instrumental in uncovering aspects of chromatin interactions and dynamics, but are limited in scale, i.e. only one or a few specific loci can be studied at one time. In addition, when whole genome dynamics is analyzed microscopically (e.g. (Zidovska et al., 2013)), positions of specific sequences have not as yet been determined.

New approaches are required to identify and quantify the molecular processes and biophysical forces that drive chromosome and nuclear compartmentalization and to characterize the dynamics and mobility of loci within these compartments. Here we describe liquid chromatin Hi-C, a Hi-C variant that quantifies the stability of chromosome conformation and chromatin interactions between loci genome-wide. We find that different types of nuclear sub-compartments differ in stability of chromatin interactions. The results suggest that compartmentalization is mainly due to strong and stable heterochromatic interactions, while associations between open regions at and around nuclear speckles, and between loci in the B1 sub-compartment enriched in polycomb binding, are more dynamic.

## RESULTS

### Measuring the stability of chromatin interactions that maintain nuclear compartmentalization

The formation of spatially segregated heterochromatic and euchromatic domains can be viewed as microphase separation of a polymer composed of different types of monomers (loci). For instance, a “block copolymer” is a polymer that contains a series of alternating blocks (e.g., A-type and B-type, or blocks of euchromatin and heterochromatin), each composed of multiple monomers (A monomers and B monomers; Figure 1A). When As attract As and Bs attract Bs, such polymer can fold into spatially segregated domains of As and Bs (Figure 1A; (de Gennes, 1979; Leibler, 1980; Matsen and Schick, 1994)). A key point is that the A and B blocks of a copolymer cannot macroscopically phase separate because they are connected: copolymers undergo microphase separation, forming microdomains of sizes comparable to the polymer coil size of the individual blocks. Applied to chromatin *in vivo*, microphase separation may underlie the formation of chromatin interaction domains. However, the biophysical forces and interaction dynamics that determine chromosome compartmentalization in cells are not known.

**Figure 1:**
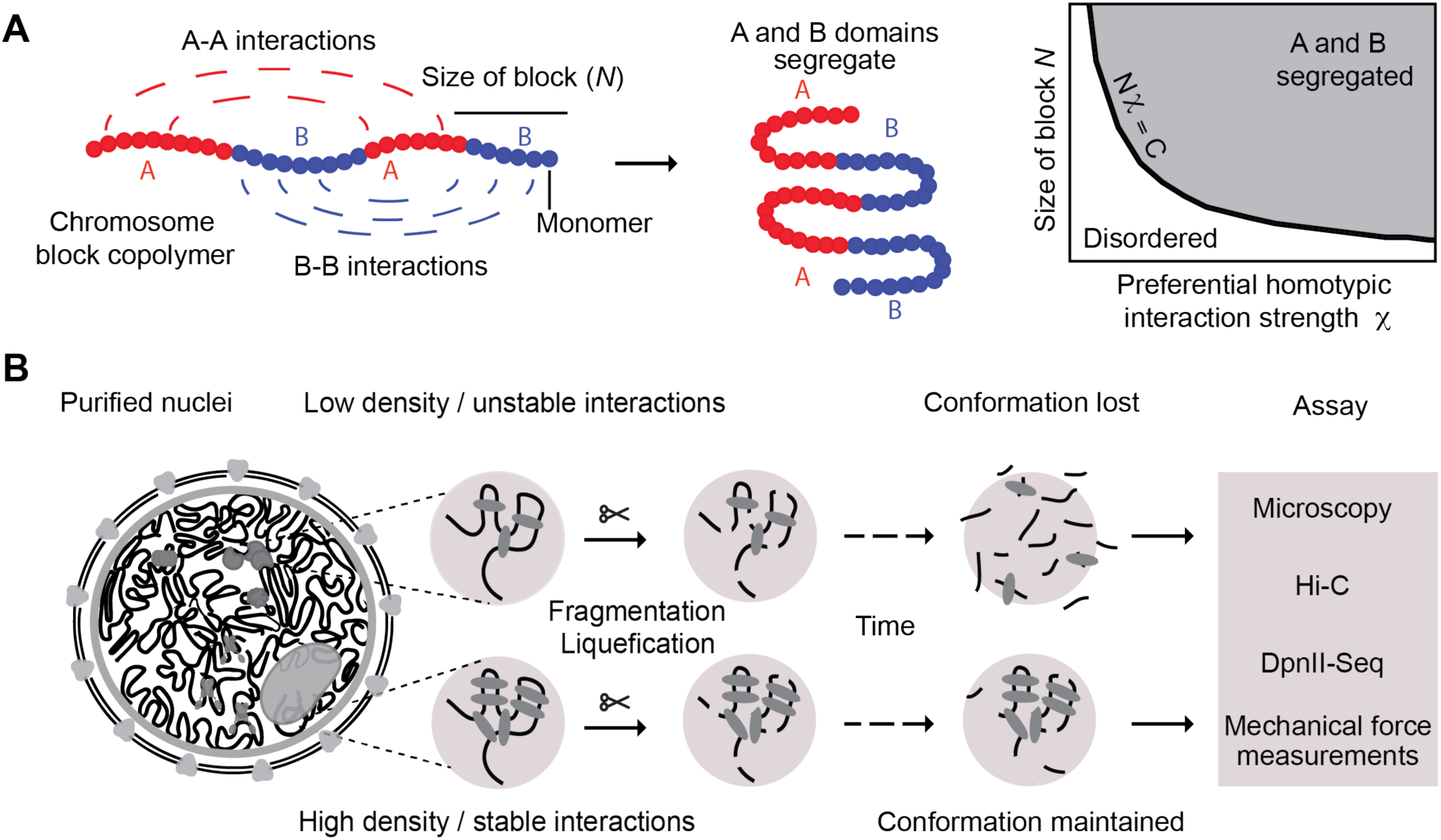
Approach for measuring chromatin interaction stability. **A**: Block copolymer composed of a series of alternating A and B blocks, each composed of a number of monomers (left). The polymer can fold into spatially segregated domains of As and Bs (middle). Flory-Huggins polymer theory predicts that spatial segregation will occur when the product of the length of the blocks *N* (the number of monomers that make up blocks) and their effective preferential homotypic interaction strength χ (difference in the strength of homotypic interactions as compared to heterotypic (A-B) interactions) is larger than a critical value *C*. **B**: Workflow to determine the stability of chromatin interactions genome-wide, DNA (black), varying chromatin features or proteins maintaining DNA conformation (grey ovals).

Whether microphase separation of a block copolymer occurs depends on the interaction strengths between monomers as well as the lengths of the blocks of monomers of each type (Figure 1A). Flory-Huggins polymer theory predicts that spatial segregation will occur when the product of the length of the blocks (*N*, the number of monomers that make up blocks) and their effective preferential homotypic interaction strength (χ, a parameter that represents the difference in the strength of homotypic interactions as compared to heterotypic (A-B) interactions) is larger than a critical value *C* (de Gennes, 1979; Leibler, 1980; Matsen and Schick, 1994); see Supplemental Materials). Large blocks of a polymer can spatially segregate even when attractive interactions among monomers are weak, while short blocks will only phase separate when interactions are sufficiently strong. Given that in mammals, domains of heterochromatin and euchromatin are typically large (hundreds of kb up to several Mb), spatial segregation can occur even when attractive forces between monomers (loci) are weak. Thus, a wide range of interaction strengths between loci within A and B compartments could lead to a spatially segregated genome. The corollary is that observation of spatially compartmentalized chromosomes per se, e.g., by Hi-C, does not provide quantitative information about the strength of the interactions that drive this organization. Should quantitative measurements of chromatin interaction strength or stability be available genome-wide, such data would allow deeper molecular understanding of the mechanisms leading to chromosome and nuclear compartmentalization, by relating these measurements to binding of specific factors, association with sub-nuclear structures, and the presence of chromatin modifications and processes such as transcription and replication.

The dependence of microdomain formation on the product of block size and interaction strength suggests an experimental approach to quantify the strengths and dynamics of interactions between individual loci that drive chromosome compartmentalization (Figure 1A, B). One can start with a compartmentalized state of the genome and fragment the chromosomes, e.g., by *in situ* restriction digestion, and then identify conditions where chromatin fragments become so short that the chromatin interaction strength between the segments is not sufficient to maintain a phase- or microphase-separated state. As a result, chromosomal domains and compartments will disassemble over time and the chromosomal fragments of different type (e.g., As and Bs) will become mixed, i.e. chromatin becomes liquid-like as opposed to a glassy state for intact chromosomes (Shi et al., 2018). The kinetics of this dissolution and mixing process can then be assessed genome-wide by Hi-C at different times after chromatin fragmentation. Domains formed by strong, stable, and abundant interactions will dissociate more slowly than domains formed by weak, unstable, or infrequent interactions (Figure 1A, B). Such approach will identify the minimum length of the blocks of monomers required for phase separation, the strength and stability of chromatin interactions, the dissolution kinetics of initially phase-separated sub-nuclear domains upon fragmentation, and how these parameters vary along the genome. Here we describe such strategy that we call liquid chromatin Hi-C.

### Chromosome conformation in isolated nuclei

To facilitate enzymatic fragmentation of chromosomes, we isolated nuclei from K562 cells, a cell line with extensive public data on chromosome conformation and chromatin state through efforts of the ENCODE project (ENCODE-Project-Consortium, 2012). We performed four analyses to demonstrate that chromosome conformation in isolated K562 nuclei was the same as that in intact cells. First, DAPI staining and imaging showed that nuclei were intact with Lamin A as a ring at the nuclear periphery (Figure 2A). Second, we performed 3C (Dekker et al., 2002) to assess known looping interactions between the beta-globin locus control region and the expressed gamma-globin genes (Chien et al., 2011; Dostie et al., 2006). These interactions were readily detected in purified nuclei, as they were in intact cells (Figure S1). Third, 5C analysis (Dostie et al., 2006) of a 1 Mb region surrounding the beta-globin locus showed that known CTCF-mediated interactions were also preserved (Figure S1; (Dostie et al., 2006; Kang et al., 2017; Splinter et al., 2006; Tolhuis et al., 2002). Fourth, genome-wide Hi-C analysis (Belaghzal et al., 2017; Lieberman-Aiden et al., 2009) confirmed that chromosome territories, compartments (determined by principle component analysis, with compartments captured by the first principle component (PC1 (Belaghzal et al., 2017; Lieberman-Aiden et al., 2009)), TADs, and CTCF-CTCF loops were intact in isolated nuclei and quantitatively similar to those in intact cells (Figure S1). We conclude that chromosome conformation and nuclear compartmentalization as detected by 3C-based assays are intact in purified nuclei.

**Figure 2:**
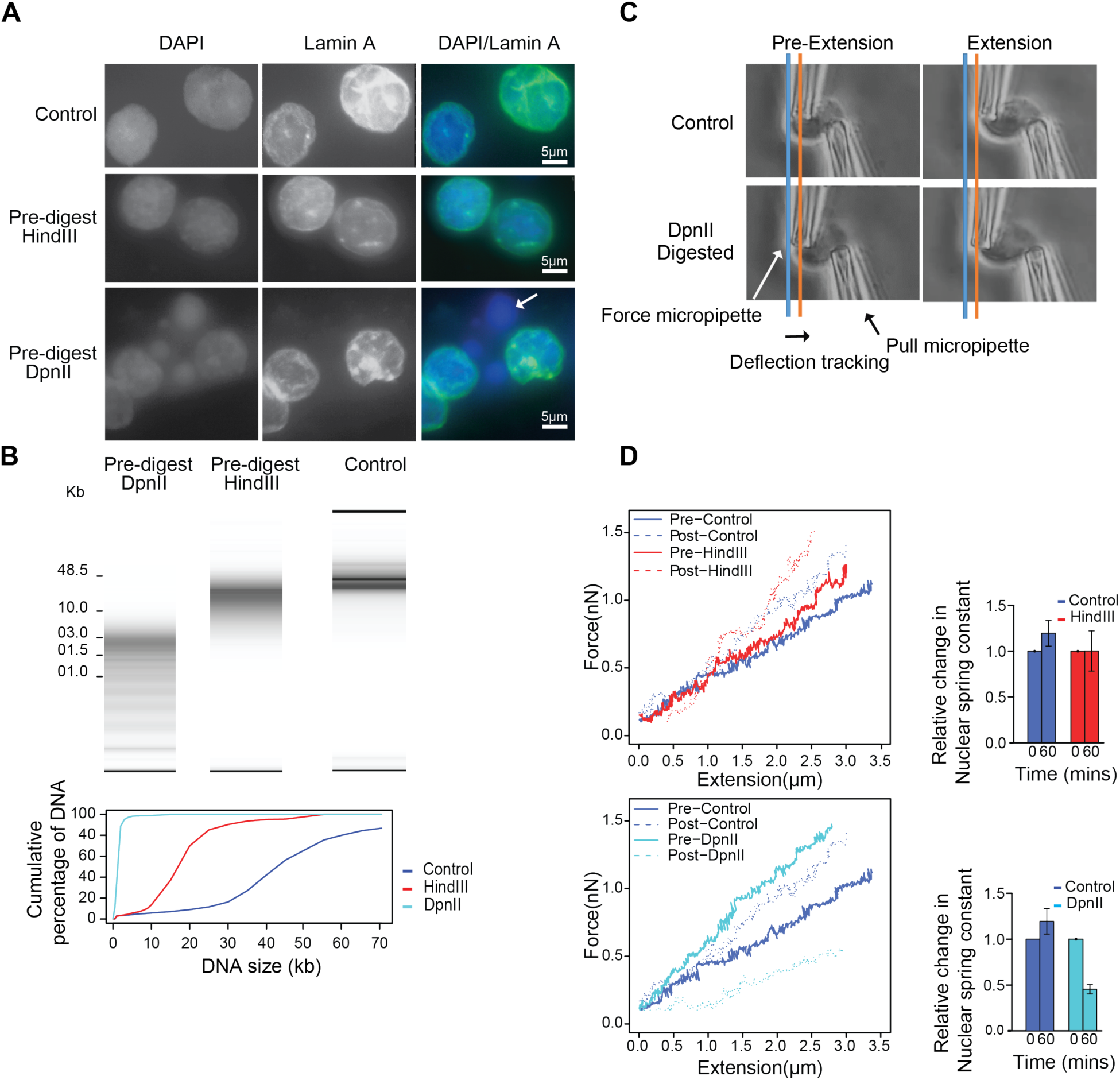
Extensive fragmentation of chromatin leads to liquefied chromatin. **A**: Nuclear and chromatin morphology before and after chromatin fragmentation. Top row: control nuclei in restriction buffer, middle row nuclei digested for 4 hours with HindIII. Bottom row: nuclei digested for 4 hours with DpnII. Nuclei were stained with DAPI (left column), with antibodies against Lamin A (middle column). The right column shows the overlay of the DAPI and Lamin A stained images. HindIII digestion did not lead to major alteration in nuclear morphology and chromatin appearance, while DpnII digestion led to the appearance of DAPI stained droplets (arrow) exiting the nuclei. **B**: Top: DNA purified from undigested nuclei, and nuclei pre-digested with DpnII and HindIII was run on a Fragment Analyzer. Bottom: cumulative DNA length distributions calculated from the Fragment Analyzer data. **C**: Micromanipulation of single nuclei. Isolated nuclei were attached to two micropipettes at opposite ends. Nuclei were extended by moving the right micropipette (Extension micropipette) and the force required was calculated from the deflection of the calibrated “force” (left) pipette. Blue and orange lines indicate the position of the force pipette before and after extension for control nuclei. After digestion of nuclei with DpnII (bottom) extension required less force as indicated by the much smaller deflection of the force pipette as compared to control nuclei (see also Supplemental Movies 1 and 2). **D.** Force-extension plots (left) for control nuclei before and 60 minutes after incubation in restriction buffer (pre- and post control), for nuclei before and after digestion with DpnII, and for nuclei before and after HindIII digestion. Right panel: relative change in nuclear spring constants, calculated from the slopes of the force-extension plots shown on the left. Bars indicate standard error of the mean (n = 5 DpnII pre-digested nuclei, and n = 4 HindIII pre-digested nuclei).

### Extensive chromatin fragmentation leads to the formation of liquid chromatin

Next, we determined the effect of different levels of chromatin fragmentation on overall nuclear organization. We incubated purified nuclei for four hours with restriction enzymes that digest chromatin with different frequencies. To quantify the extent of digestion, DNA was purified, and the size distribution of DNA fragments was determined using an Agilent Fragment Analyzer. Digestion with HindIII resulted in fragments that ranged in size from ∼10-25 kb (Figure 2B). A minority of molecules was over 25 kb (<15%), indicating that most of the genome was fragmented to a similar extent. Digestion with DpnII resulted in fragments that ranged in size between ∼1 and ∼6 kb, with less than 6% of fragments >6 kb (Figure 2B). Microscopic inspection of nuclear morphology by DAPI and Lamin A immunofluorescence staining showed that fragmentation of chromatin with HindIII had only minor effects on nuclear morphology (Figure 2A). We did occasionally notice some small amounts of DNA emerging as tiny droplets from the nuclear periphery, suggesting that HindIII digestion led to some solubilization of chromatin. In contrast, fragmentation of chromatin with DpnII led to large-scale alteration of nuclear morphology as detected by DAPI staining, and large droplets of apparently liquid chromatin (not surrounded by Lamin A) emerged from the nuclear periphery (Figure 2A, arrow).

We next tested whether different chromatin fragmentation levels had an effect on nuclear stiffness, which reflects the integrity of chromosome conformation and chromatin interactions inside the nucleus. We previously showed that single-nucleus isolation and whole-nucleus extension via micromanipulation with micropipettes provides a reliable and robust way to measure the stiffness of the nucleus (Stephens et al., 2017). K562 cells were pre-treated with the actin depolymerizing drug latrunculin A to allow isolation of the nucleus. Isolated nuclei were attached to two micropipettes at opposite ends, and the whole nucleus was extended by moving an extension micropipette. The deflection of a force micropipette multiplied by its premeasured spring constant provides a measure of the force (Figure 2C; Supplemental Movie 1, 2). This data provides a force vs. extension plot (Figure 2D, plots on the left), in which the slope of the line fitted to the data is the nuclear spring constant in nN/μm (Figure 2D, bar plots on the right). Extension between 0 – 30% strain measures the chromatin-dominated regime of nuclear force response (Stephens et al., 2017; Stephens et al., 2018), and simulations of nuclear mechanics suggest chromosomal interactions contribute to this regime (Banigan et al., 2017). Isolated single nuclei were measured for their native spring constant before treatment with a restriction endonuclease and then re-measured 60 minutes post-treatment. Nuclei treated with control conditions (only restriction buffer added to the media) showed a slight stiffening of the nucleus (Figure 2D). Treatment of nuclei with HindIII did not significantly decrease the stiffness compared to controls. In sharp contrast, DpnII-treated nuclei displayed a significant decrease (∼75%) in stiffness, consistent with previous experiments treating nuclei isolated from mouse embryonic fibroblasts with AluI (Stephens et al., 2017). We conclude that global chromosome and nuclear organization can tolerate genome-wide fragmentation to 10-25 kb segments, indicating that sufficient numbers of relatively stable chromatin interactions are maintained between these large fragments throughout the genome to maintain nuclear stiffness. In contrast, fragmenting the genome to smaller than 6 kb segments results in extensive loss of chromatin morphology, loss of chromatin-mediated stiffness, and the appearance of a liquid-like state of chromatin.

### Liquid chromatin Hi-C analysis reveals that compartmental segregation requires chromatin fragments larger than 6 kb

To determine how chromosome folding and nuclear compartmentalization across the genome is altered as a function of chromatin fragmentation level, we applied Hi-C before (conventional Hi-and after chromatin liquefication (liquid chromatin Hi-C). Nuclei were digested with either HindIII or DpnII for 4 hours followed by formaldehyde fixation and Hi-C analysis (Figure S2A). Liquid chromatin Hi-C interaction maps obtained from nuclei that were pre-digested with HindIII were remarkably similar to those obtained with nuclei that were not pre-digested (Figure 3A). The relationship between interaction frequency and genomic distance was largely unaffected, with only a slight redistribution to longer-range interactions (Figure 3B). The ratio of intra-vs. inter-chromosomal interactions was also highly similar to that in untreated nuclei (Figure 3B). Compartments (Figure 3D) were readily detectable as a plaid pattern in Hi-C interaction maps. This pattern is captured by the first principle component (PC1). Compartment positions were unaffected (Spearman ρ= 0.99).

**Figure 3:**
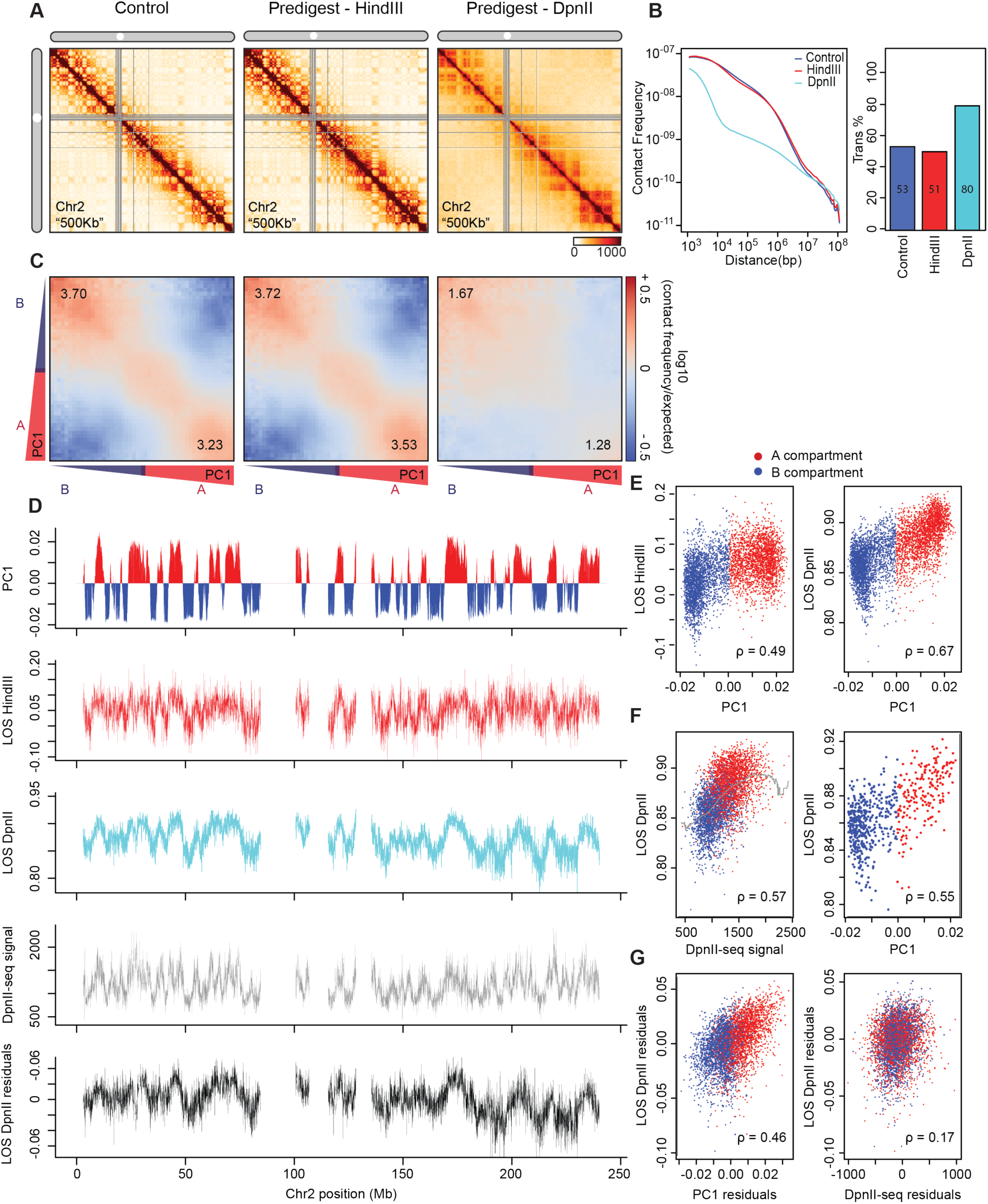
Hi-C analysis reveals chromosome disassembly upon chromatin liquefication. **(A)** Hi-C interaction maps of chromosome 2 binned at 500 kb. Left: interaction map for control nuclei in restriction buffer prior to pre-digestion. Middle: nuclei pre-digested for 4 hours with HindIII prior to Hi-C. Right: nuclei digested for 4 hours with DpnII prior to Hi-C (see Figure S2A). **(B)** Left: genome-wide interaction frequency as function of genomic distance for control nuclei (dark blue), nuclei pre-digested with HindIII (red), and nuclei pre-digested with DpnII (cyan). Right: percentage of inter-chromosomal (trans) interaction frequencies. **(C)** Compartmentalization saddle plots: average intra-chromosomal interaction frequencies between 40 kb bins, normalized by expected interaction frequency based on genomic distance. Bins are sorted by their PC1 value derived from Hi-C data obtained with control nuclei. In these plots preferential B-B interactions are in the upper left corner, and preferential A-A interactions are in the lower right corner. Numbers in corners represent the strength of AA interactions as compared to AB interaction and BB interactions over BA interactions (Figure S4B). **(D)** Top plot: Eigenvector 1 values (PC1, 40 kb resolution) across a section of chromosome 2, representing A (red) and B (blue) compartments. Second plot: Loss of pair-wise interactions “LOS” (Methods and Figure S2B) along chromosome 2 at 40 kb resolution for nuclei pre-digested with HindIII. Third plot: LOS for nuclei pre-digested with DpnII. Fourth plot: DpnII-seq signal along chromosome 2 at 40 kb resolution. Bottom plot: LOS-residuals for nuclei pre-digested with DpnII after correction for DpnII signal. **E)** Correlation between LOS for nuclei pre-digested with HindIII (left) or DpnII (right) and PC1 (for chromosome 2, Spearman correlation values are indicated). **F)** Left: correlation between LOS for nuclei pre-digested with DpnII and DpnII-seq signal (for chromosome 2). Grey line indicates moving average used for residual calculation. Right: correlation between LOS for nuclei pre-digested with DpnII and PC1 for loci cut to the same extent by DpnII (1000-1100 DpnII-seq reads/ 40 kb bin; for chromosome 2). Spearman correlation values are indicated. **G)** Left: partial correlation between residuals of LOS for nuclei pre-digested with DpnII and residuals of PC1 after correcting for correlations between LOS and DpnII-seq and PC1 and DpnII-seq signal. Right: partial correlation between residuals of LOS for nuclei pre-digested with DpnII and residuals of DpnII-seq signal after correcting for correlations between LOS and PC1 and DpnII-seq signal and PC1. Spearman correlation values are indicated.

Chromosome compartment strength can be visualized and quantified by plotting interaction frequencies between pairs of 40 kb loci arranged by their values along the first eigenvector (PC1) to obtain compartmentalization saddle plots (Figure 3C). In these plots the upper left quadrant represents B-B interactions and the lower right corner represents A-A interactions. Interestingly, in nuclei pre-digested with HindIII, the strength of preferential A-A and B-B interactions (Figure 3C; Supplemental Figure 2C shows a replicate) was somewhat increased, indicating stronger segregation of A and B compartments. This observation, puzzling at first, is in fact readily understood when chromosomes fold as block co-polymers. Polymer theory predicts that very mild fragmentation of a copolymer can enhance phase (or microphase) separation by removing covalent linkages between A and B blocks as long as the fragments are still large enough to sufficiently attract each other (See Methods). Our results show that chromatin fragments of 10-25 kb are long enough to allow stable segregation of A and B compartments genome-wide.

Much more extensive changes in chromosome conformation were observed when nuclei were pre-digested for 4 hours with the frequent cutting enzyme DpnII (Figure 3A) followed by formaldehyde fixation and Hi-C analysis. We observed a considerable loss in shorter range (<10 Mb) intra-chromosomal interactions, with a gain of longer range (>10 Mb) interactions and inter-chromosomal interactions (Figure 3B). The gain in inter-chromosomal interactions appeared to be the result of random mixing of As and Bs as the preference for interchromosomal A-A and B-B interactions decreased. Moreover, compartment strength *in cis* was greatly reduced with a greater relative reduction evident in the A compartment (Figure 3C). This more prominent loss of A-A interactions compared to B-B interactions is also apparent from direct visual inspection of the Hi-C interaction maps: while in Hi-C maps from undigested nuclei a plaid interaction pattern is clearly visible with two alternating patterns of chromatin interactions (representing A and B compartments), in liquid chromatin Hi-C maps from DpnII-pre-digested nuclei one of these two patterns is weakened more (corresponding to the A compartment). Combined, these observations show that fragmentation in <6 kb fragments leads to loss of spatial segregation of A and B compartments and dissolution of chromosome conformation genome-wide but with more extensive loss of the A compartment.

### Quantification of chromosome conformation dissolution upon chromatin fragmentation

Loss of chromosome conformation and dissolution of chromosomal compartments will result in random mixing of previously spatially separated loci both *in cis* and *in trans*. In Hi-C this will be apparent by loss of short-range interactions and gain in longer range and inter-chromosomal interactions. We used this phenomenon to quantify for each locus along the genome the extent of loss of chromosome conformation upon chromatin fragmentation. Specifically, using Hi-C data binned at 40 kb resolution we developed a metric which represents the percentage change in short range intra-chromosomal interactions (up to 6 Mb) for each fragmentation condition relative to control nuclei, which we call loss of structure (LOS) (Figure S2B).

We first calculated LOS after 4 hours for chromatin fragmented with HindIII. We observe that in general short-range interactions are only somewhat reduced, consistent with the above observation that interactions are somewhat redistributed to longer-range interactions (Figure 3B). When LOS is plotted along chromosomes (Figure 3D), we observed that LOS is not uniform: some regions display more loss of short-range interactions than others. To determine how the loss of interactions is related to A and B compartments, we compared LOS to the PC1 value that captures compartmentalization (Lieberman-Aiden et al., 2009) and observed that LOS was positively correlated with PC1 (Figure 3D, 3E left panel): ∼3 - 12% loss for loci in A and <5% loss for loci in B (Figure 3E). We note that this effect is very small and close to technical variation between replicates. Thus, chromosome conformation and compartmentalization are largely intact in nuclei pre-digested with HindIII (See Supplemental Figure S2C for a replicate).

We performed the same analysis for nuclei pre-digested with DpnII for 4 hours followed by formaldehyde fixation and Hi-C. Consistent with the micromechanical measurements described above, we find an extensive loss of chromosome conformation, with LOS generally >80%. LOS varies along chromosomes and is strongly positively correlated with PC1 with loci in the A compartment displaying the largest loss (Figure 3D, 3E). These results again show that chromatin fragmentation to <6 kb fragments leads to extensive genome-wide dissolution of chromosome conformation, random mixing of loci, and loss of spatial segregation of A and B compartments, with the A compartment affected to the largest extent.

### Independent contributions of compartment status and fragmentation level to chromatin dissolution

As outlined above, phase segregation of polymers is predicted to depend on both the length of fragments and the attractive forces between them. Therefore, one explanation for the greater effect of fragmentation on chromatin interactions and chromosome conformation in the A compartment could be that DpnII cuts more frequently in the open and potentially more accessible A compartment. To assess this, we determined the cutting frequency of DpnII in isolated nuclei across the genome (DpnII-seq; Supplemental Figure S3). Nuclei were digested with DpnII for 4 hours, and free ends were filled in with biotinylated nucleotides. After shearing, biotin-containing fragments were isolated, DNA was sequenced and reads were mapped to the genome. The frequency of fragments mapping with one end at a DpnII restriction site was calculated along chromosomes at 40 kb resolution and compared to PC1. We find that digestion frequency is positively correlated with PC1 and with LOS (Figure 3D, 3F left panel; Figure S2D).

To determine whether the correlation between LOS and PC1 is only due to the fact that DpnII digestion is correlated with PC1 we calculated the partial correlation between LOS and PC1 after correcting for the correlations of PC1 and LOS with DpnII digestion frequency (see Methods for details). We find that the residuals of PC1 and LOS are still highly correlated (Spearman ρ = 0.46 for chromosome 2; Figure 3G). To illustrate the correlation between LOS and PC1 independent of fragmentation level directly we selected a set of loci along chromosome 2 that are all cut to the same extent (1000-1100 reads in the DpnII-seq dataset).

When we plot LOS vs. PC1 for this set we find a strong correlation (Figure 3F right panel, Spearman ρ = 0.55). We also determined the partial correlation between LOS and DpnII-seq signal after correcting for their correlations with PC1. We find that after this correction LOS and DpnII-seq signal remain correlated (Spearman ρ = 0.17, Figure 3G right panel). We conclude that when generally digested to <6 kb fragments both compartment status (PC1) and fragment size contribute to LOS. Importantly, this implies that the interaction strength between chromatin segments is related to their compartment status.

### Dissociation kinetics of chromatin interactions and compartments

The observation that pre-digestion of chromatin with DpnII produces chromatin fragments that are too small to maintain segregation of chromatin in A and B compartments allowed us to measure the dissociation kinetics of compartments and stability of chromatin interactions as loci become mobile and start to mix. We first determined the kinetics of chromatin fragmentation (Figure S4A). We digested nuclei with DpnII for different amounts of time, ranging from 5 minutes to 16 hours. At each time point, we isolated DNA to determine the extent of digestion (Figure 4A). After 5 minutes the size range of fragments was between 3 and 15 kb (80% of fragments). After one hour 80% of DNA fragments were smaller than 7 kb and during the subsequent hours of digestion the fragments became gradually shorter. After 16 hours of digestion 85% of fragments were smaller than 3.5 kb. We again sequenced DNA ends to determine the distribution of DpnII cuts across the genome (Figure 4B). We find that at all timepoints the number of DpnII cuts per 40 kb bin was correlated with PC1 (Figure 4B), but that the pattern did not change much over time (Figure 4B, correlation matrix; Figure S5). We conclude that digestion is more frequent in the A compartment at all timepoints, but that the pattern of fragmentation along chromosomes is relatively stable over the timecourse.

**Figure 4:**
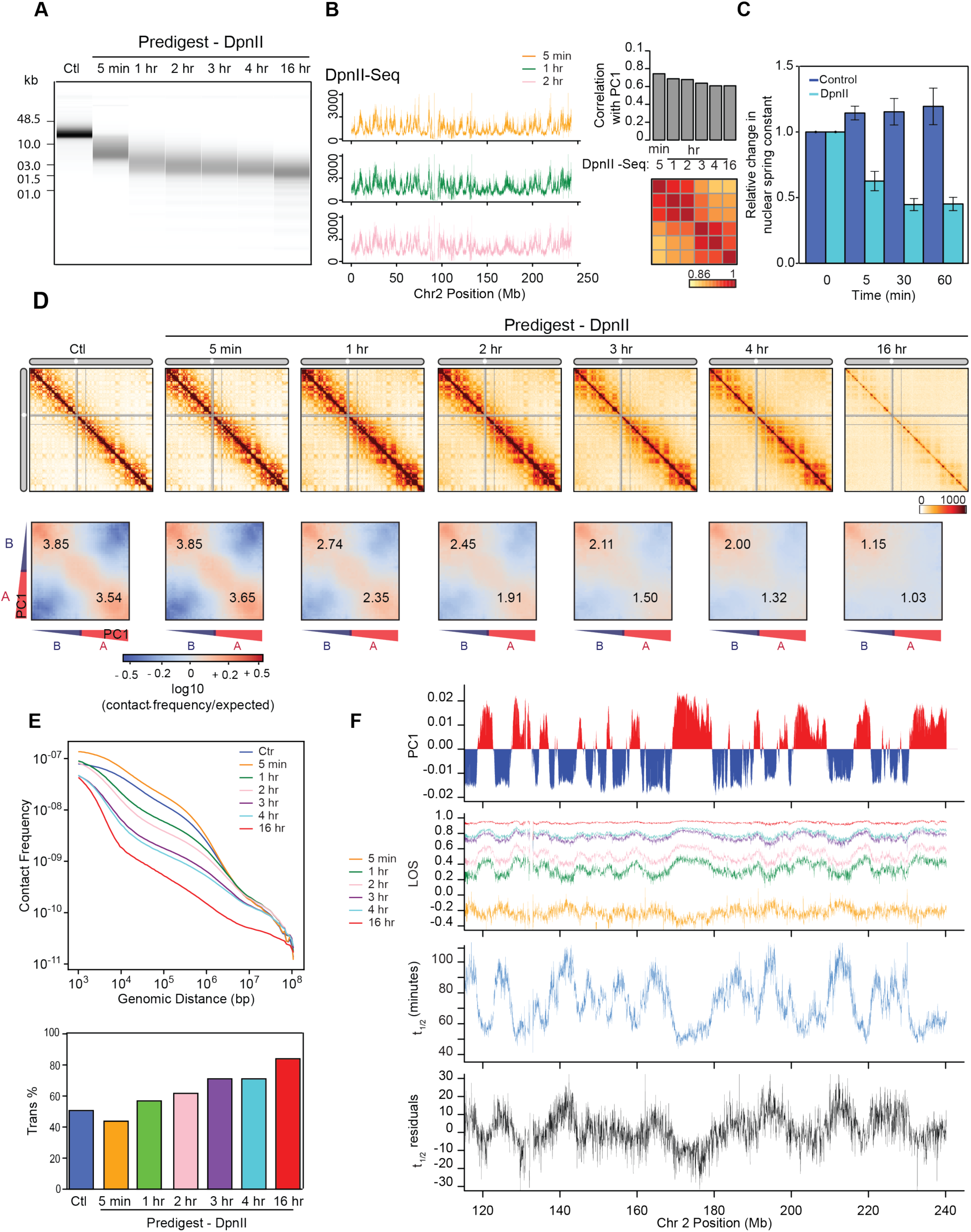
Kinetics of chromatin fragmentation and chromatin dissolution. **(A)** DNA purified from undigested nuclei, and nuclei pre-digested with DpnII for different time points were run on a Fragment Analyzer. **(B)** Left: DpnII-seq signals along chromosome 2 binned at 40 kb resolution after digestion for 5 minutes, 1 hour and 2 hours. Right: correlation between DpnII-seq signals and PC1 and between DpnII-seq signals at different time points. **(C)** Relative change in nuclear spring constant (nN/µm) after DpnII fragmentation at different time points. Spring constant is significantly decreased after 5 minutes and at background level by 1 hour (p = 0.002, two-tailed t-test). **(D)** Top row: Hi-C interaction maps of chromosome 2 binned at 500 kb. Control: nuclei in restriction buffer for 4 hours. Pre-digest DpnII: nuclei were pre-digested with DpnII for 5 minutes up to 16 hours. (Figure S4A). Bottom row: compartmentalization saddle plots for the corresponding conditions. Numbers indicate strength of A-A and B-B interactions. **(E)** Top: genome-wide interaction frequency as function of genomic distance for Hi-C data shown in panel (D). Bottom: percentage of inter-chromosomal (trans) interactions genome-wide for control nuclei and for nuclei pre-digested with DpnII for up to 16 hours. **(F)** Top: PC1 along a section of 120 Mb of chromosome 2. Second plot: LOS along chromosome 2 at 40 kb resolution for all time points (Figure S2B). Third plot: half-life (t_1/2_) values derived from the exponential fit of the six time-points for every 40 kb bin (Figure S4C). Bottom plot: residuals of t_1/2_ after correcting for correlations between t_1/2_ and DpnII-seq (DpnII-seq data for t = 1 hour).

Micromanipulation was again used to measure the nuclear spring constant corresponding to nuclear stiffness. Nuclei displayed a significant loss in stiffness within 5 minutes, reaching background levels after 1 hour ((Stephens et al., 2017), Figure 4C) when chromatin fragments are smaller than 7 kb. Combined these analyses show that the bulk of DNA fragmentation and chromatin liquefication occurs within the first hour, after which the nuclei have completely lost their nuclear stiffness.

Next, we used liquid chromatin Hi-C to determine the kinetics with which chromosome conformation and compartmentalization are lost after chromatin fragmentation. Nuclei were pre-digested with DpnII for 5 minutes up to 16 hours, followed by formaldehyde fixation and Hi-C analysis (Figure S4A). Interestingly, after 5 minutes of pre-digestion chromosome conformation and compartmentalization are intact, even though chromatin was fragmented to 3-15 kb segments before fixation and nuclear stiffness was significantly reduced (Figure 4C, D). Further, the percentage of intra-chromosomal interactions especially for loci separated by <1 Mb was increased (Figure 4E). Compartmentalization is somewhat increased for the A compartment (Figure 4D). These increases may be due to the fact that fragments are on average around 10 kb long, only somewhat shorter than after HindIII digestion, but longer than after 4 hours of DpnII digestion (Figures 2, 3). As discussed above mild fragmentation of a block copolymer is predicted to enhance both microphase or phase separation, depending on the cut frequency (see Methods). The results also indicate that interactions among fragments of around 10 kb are strong enough for compartmentalization to occur.

At subsequent time points, when most chromatin fragments are <7 kb long and nuclear stiffness is completely lost, we observe increased loss of intra-chromosomal interactions and concomitant increased inter-chromosomal interactions genome-wide (Figure 4D, 4E). Compartmentalization, as quantified by the preference of A-A and B-B interactions over A-B interactions, is progressively lost (Figure 4D, lower row of heatmaps, Figure S4B). A-A interactions disappear faster than B-B interactions. After 16 hours, only a low level of preferential B-B interaction remains. This analysis shows that chromatin fragmentation transiently leads to more frequent intra-chromosomal interactions and stronger compartmentalization due to partial digestion which is then followed by progressive genome-wide dissolution of chromosome conformation as fragments become too short, and interactions between these fragments too weak to maintain compartmentalization.

### Quantification of the half-life of chromosome conformation across the genome

To quantify the kinetics of loss of chromosome conformation and compartmentalization, we calculated LOS genome-wide for each time point (Figure 4F). At t = 5 minutes, LOS is generally negative indicating a gain in chromatin interactions: on average ∼25% gain of intra-chromosomal interactions between loci separated by <6 Mb, consistent with the initial increase in overall intra-chromosomal interactions described above (Figure 4E). LOS is inversely correlated with PC1, indicating that loci located within A compartments gain more intra-chromosomal interactions than loci located within B compartments (Spearman ρ = −0.53 for chromosome 2, Spearman ρ = −0.49 genome-wide). At subsequent time points, LOS is increasingly positive as intra-chromosomal interactions are progressively lost and non-specific inter-chromosomal interactions are gained. LOS is the highest for loci located in the A compartment. At t = 16 hours, LOS is generally as high as 90%, intra-chromosomal interactions are low (<20% of total), and only preferential B-B interactions are still observed in the Hi-C interaction map (Figure 4D). Very similar results were obtained with an independent replicate of this time course experiment (see below). In that experiment, we again observed an initial transient gain in intra-chromosomal interactions after 5 minutes of pre-digestion, followed by a progressive loss of chromosome conformation genome-wide, but especially in the A compartment.

Now that we measured LOS as a function of time after chromatin fragmentation, we could calculate the dissociation rates of chromatin interactions genome-wide. LOS as a function of time for each 40 kb locus was fitted to an exponential curve, which was then used to calculate the time at which each locus lost 50% of its intra-chromosomal (<6 Mb) interactions (Figure S4C). We refer to this time as the half-life t_1/2_ (minutes) of chromatin interactions at each locus (Fig. 4F). A short t_1/2_ represents unstable interactions, while more stably interacting loci will have longer t_1/2_ values. Examining t_1/2_ along chromosomes, we observe a strong inverse correlation with PC1 (Spearman ρ = −0.87; Figure S4F): interactions in the A compartment dissolve relatively fast (t_1/2_ = 40-80 minutes) while interactions in the B compartment dissolve slower (t_1/2_ = 60-120 minutes; Figure S4D). We also calculated t_1/2_ genome-wide for the second independent time course experiment and find a strong correlation between t_1/2_ calculated from the two datasets (Spearman ρ = 0.78 for chromosome 2; Spearman ρ = 0.76 genome-wide; Figure S4E). The value of t_1/2_ is proportional to the dissociation rate constant and thus independent of the initial level of intra-chromosomal interactions for a given locus. Indeed, t_1/2_ remains highly correlated with PC1 even after correcting for correlations between the initial level of intra-chromosomal (<6 Mb) interactions and t_1/2_ and between the level of intra-chromosomal interactions and PC1 (Spearman ρ = −0.82, Figure S4F, G).

We can also estimate the half-life of compartmentalization directly by calculating the rate of loss of preferential A-A and B-B interactions specifically from the compartmentalization saddle plots shown in Figure 4D. Using this metric, similar values of t_1/2_ were observed: 50% loss of preferential A-A interactions was observed after ∼60 minutes, while a 50% loss of preferential B-B interactions occurred after ∼115 minutes.

### Independent contributions of compartment status and fragmentation level to the half-life of chromatin interactions

Similar to LOS, t_1/2_ is correlated with DpnII digestion frequency at all timepoints (Figure S5A). We again determined the individual contributions of compartment status and fragmentation level to t_1/2_. We calculated the partial correlation between t_1/2_ and PC1 after correcting for correlations between PC1 and t_1/2_ with DpnII cutting frequency. We find that t_1/2_ and PC1 remain strongly correlated (Figure 4F), regardless of which DpnII fragmentation dataset (genome wide Spearman ρ ranging from −0.41 to −0.60, t = 5 min up to t = 16 hours) was used for the calculation of the partial correlation (Figure S5A, S5B). This is illustrated by plotting t_1/2_ residuals as a function of DpnII fragmentation level (Figure S5C). Although loci in the A compartment are often cut more frequently than loci in the B compartment, when comparing loci cut with similar frequency, loci in the A compartments had shorter half-lives (Figure S5C). We also calculated the partial correlation between t_1/2_ and fragmentation level after correcting for their correlations with PC1 and find that fragmentation level also contributes to t_1/2_ (genome wide Spearman ρ = −0.48, chromosome 2 Spearman ρ = −0.31). We conclude that after digestion with DpnII dissolution kinetics are determined by both the compartment status of loci and their fragmentation level.

Finally, we considered whether we could have overestimated the t_1/2_ for loci in the B compartment because fragmentation of these loci could be slower than for loci in the A compartment. We reasoned that because after 1 hour incubation with DpnII digestion is largely complete, calculation of LOS using the Hi-C data at t = 1 hour as starting condition would provide an estimate of dissolution kinetics starting at a timepoint when A and B compartments are both extensively fragmented. We find that LOS, and t_1/2_ calculated this way are still strongly correlated with PC1, and this correlation remains strong after correcting for fragmentation level (Supplemental Figure S5D, S5E, S5F).

### Compartment size and boundaries influence chromatin interaction stability

We next explored the correlation between PC1 and t_1/2_ in more detail. Although these two parameters are highly correlated, visual inspection of the data suggested that the largest A compartments appeared to have the smallest t_1/2_ while the largest B compartments had the largest t_1/2_. To quantify this, we plotted t_1/2_ for each 40 kb bin as a function of the size of the compartment that the bin was located in (Figure S6D). We find that loci within small A compartments had larger t_1/2_ values than loci in larger A compartments. Analogously, loci within small B compartments had smaller t_1/2_ values than loci in larger B compartments.

One explanation for this effect is that proximity to a compartment boundary modulates the stability of loci. To analyze this, we aggregated PC1 and t_1/2_ around A-B compartment boundaries (Figure S6C). We find that while PC1 switches sharply at boundaries, t_1/2_ changes less rapidly. As a result, loci in A compartments but near a boundary have larger t_1/2_ values than expected for their PC1 value, while loci in B compartments but near a boundary have smaller t_1/2_ values than expected for their PC1 value. This boundary effect can contribute to bins in smaller compartments having larger (A compartment) or smaller (B-compartment) t_1/2_ values than loci in larger compartments.

### Dissociation kinetics of chromatin interactions at different sub-nuclear structures

A compartments can be split into A1 and A2 sub-compartments that are both characterized by open and active chromatin, with the A1 sub-compartment the most enriched in active histone modifications. The A1 sub-compartment is found in close proximity to nuclear speckles (Chen et al., 2018). Inactive chromatin can be classified into B1, B2 and B3 sub-compartments. B2 and B3 are generally inactive domains and are located near the nuclear lamina (B2 and B3) and the nucleolus (B2) (Chen et al., 2018; Quinodoz et al., 2018; Rao et al., 2014). B1 is enriched in the repressive H3K27me3 mark, which is often associated with polycomb binding. To relate sub-compartment status to chromatin dissociation rates, we compared the residuals of t_1/2_ (after correcting for fragmentation level) for loci located in the 5 sub-compartments defined for K562 cells ((Xiong and Ma, 2019); Figure 5A). We find that residual t_1/2_ varies greatly between sub-compartments: t_1/2_ (A1) ∼ t_1/2_ (B1) < t_1/2_ (A2) < t_1/2_ (B2) ∼ t_1/2_ (B3).

**Figure 5:**
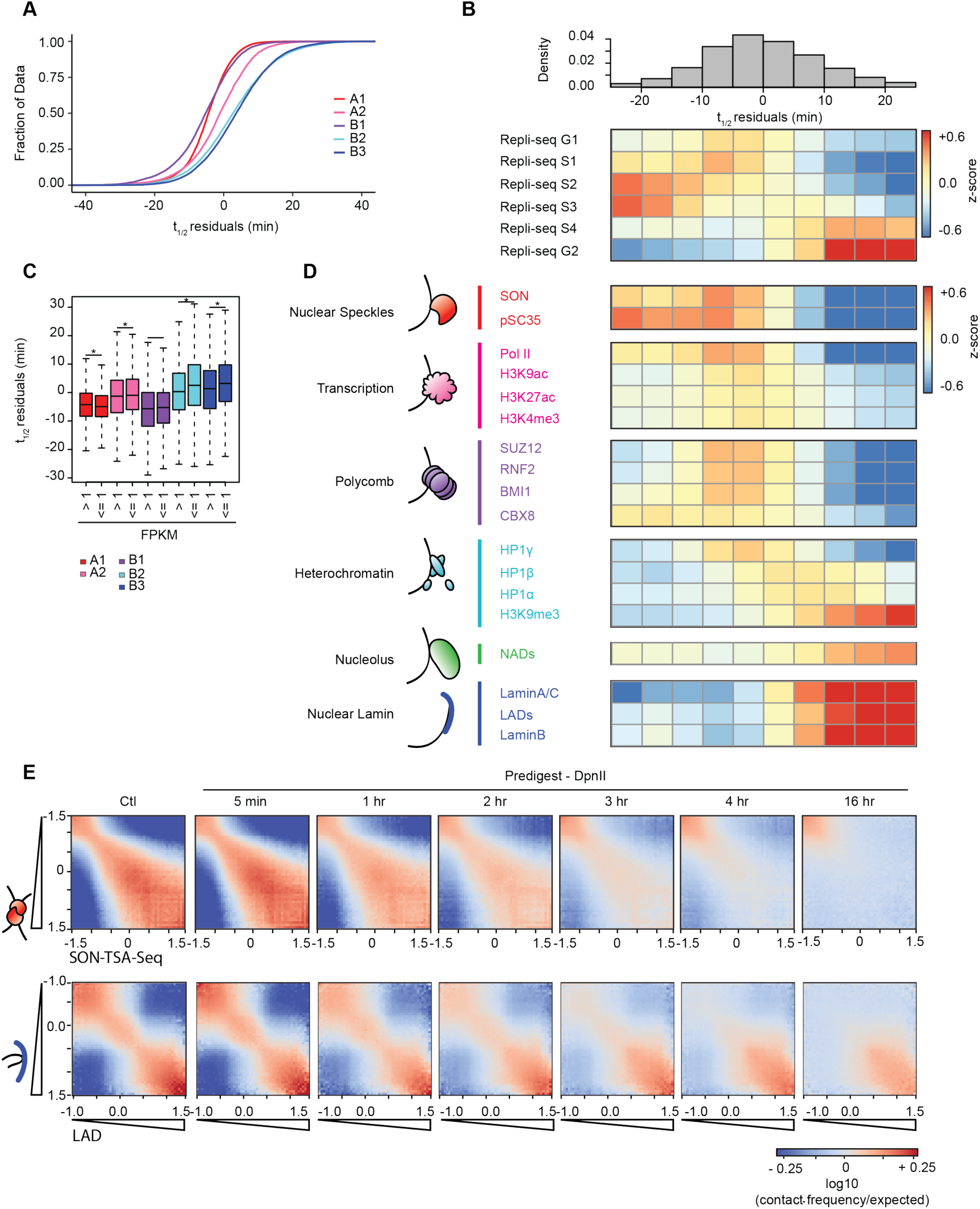
Dissociation kinetics of chromatin interactions at different sub-nuclear structures. **(A)** Cumulative distributions of residuals of t_1/2_ (in minutes) for each of the five annotated sub-compartments. **(B)** Top: the genome was split into 10 bins, where each bin corresponds to sets of loci that share the same t_1/2_ residual interval. Bottom: For each t_1/2_ residual interval the mean z-score signal of Repli-Seq data in different phases of the cell cycle G1, S1-4, G2. **(C)** Boxplot of t_1/2_ residuals for bins with expressed genes (mean FPKM > 1) and bins with low or no expression (mean FPKM <=1) stratified by sub-compartment. Significance determined by two-sample two tailed *t*-test *(p < 0.003). **(D)** Heatmap of mean z-score signal enrichment for various markers of sub-nuclear structures (See Methods) stratified by t_1/2_ residual intervals (top of panel B). For loci in each t_1/2_ residual interval the mean z-score was quantified for different chromatin features. **(E)** Homotypic interaction saddle plots for loci ranked by their association with speckles (as detected by SON-TSA-seq, top (Chen et al., 2018)) and by their association with the nuclear lamina. Preferential pair-wise interactions between loci associated with the lamina can still be observed after several hours, whereas preferential pair-wise interactions between loci associated with speckles are lost more quickly.

It is noteworthy that interaction dissociation rates for loci in the B1 sub-compartment are as high or higher (residuals of t_1/2_ as or more negative) than those for loci in the active and open A1 sub-compartment. Many B1 sub-compartments are indeed embedded within A compartments (Figure S6C, E) and a subset is found close to nuclear speckles (Chen et al., 2018). Within the B compartment, interactions between Lamin-associated loci in the B3 sub-compartments dissociate the slowest while interactions between loci in the B2 sub-compartment dissociate somewhat faster. These observations indicate that loci associated with different sub-nuclear structures display a range of interaction stabilities. We also noted that A2 sub-compartments are frequently found near A-B boundaries (Figure S6C) and that A2 and B2 sub-compartments tend to be located in smaller A and B compartments respectively (Figure S6E), providing an additional explanation for the compartment size effect on t_1/2_ described above.

Consistent with the relation between chromatin state, sub-compartment status, and DNA replication timing (Rao et al., 2014), we find a strong correlation between t_1/2_ residuals and replication timing. We split the genome into 10 bins, where each bin corresponds to sets of loci that share the same t_1/2_ residual interval. We then explored the enrichment for varying chromatin features for each t_1/2_ residual interval (Figure 5B, D). Chromatin interactions for early replicating domains had short half-lives, while interactions for loci in later replicating domains were more stable (Figure 5B). Interestingly, loci with the shortest t_1/2_ residuals replicate in the middle of S-phase.

The strong correlation between active chromatin and relatively unstable chromatin interactions led us to examine the role of transcription in more detail (Figure 5C). For each sub-compartment, we split loci into expressed (FPKM>=1) or not expressed (FPKM<1) categories. We find that sub-compartment status is the major determinant of chromatin interaction stability, irrespective of transcriptional status. However, transcriptional status modulates t_1/2_ to some extent: in general, loci located in B2 and B3 sub-compartments are engaged in relatively stable chromatin interactions, but interactions that involve loci that are expressed have shorter half-lives. Conversely, the expression status of loci located in the A1, A2 sub-compartments had only very minor effect on the residual t_1/2_. Expressed loci in A1 or A2 sub-compartments had slightly longer and shorter residual t_1/2_ values. Although statistically significant, the average difference was very small (A1 difference = 0.87 minutes; A2 difference = 0.68 minutes).

To explore the relationship between chromatin interaction stability, chromatin state, and association at and around sub-nuclear structures in more detail, we leveraged the wealth of chromatin state data available for K562 cells (Figure 5D, Figure S6A, B). Chen et al. mapped loci in K562 cells that are localized near nuclear speckles using the recently developed TSA-seq method (Chen et al., 2018). We find that loci near the speckle-associated protein pSC35 are engaged in the most unstable interactions in the genome. A similar result was obtained for an independent TSA-seq dataset for the speckle associated protein SON. Similarly, transcriptionally active loci, identified by ChIP-seq for a range of histone modifications and factors associated with open chromatin such as H3K4me3 and RNA PolII, were also involved in relatively unstable chromatin interactions, though for some marks the residual t_1/2_ values varied more widely.

Interestingly, interactions for loci bound by polycomb complexes (a subset of which are in the B1 sub-compartment) were as unstable as active and speckle associated loci (Figure 5D, S6B). This suggests that polycomb-bound domains, are held together by highly dynamic interactions. Interestingly, half-lives differed for loci bound by different polycomb subunits. Loci with the shortest t_1/2_ values are enriched specifically for binding the CBX8 subunit. An example of a large polycomb-bound domain in K562 cells is the HoxD cluster. The cluster is around 100 kb in size and covered by the polycomb subunits Suz12, RNF2, CBX8 and BMI1 and the histone modification H3K27me3 (Figure S6G). The half-life of chromatin interactions for loci in the HoxD cluster is relatively short.

Silent and closed chromatin loci around the nucleolus or at the nuclear lamina were generally engaged in the most stable interactions (Figure 5D). Chromatin interaction stabilities for loci associated with the three distinct heterochromatin proteins 1 (HP1) differed: chromatin interactions for loci associated with HP1γ (CBX3) were relatively unstable while interactions for loci associated with HP1β (CBX1) or HP1α (CBX5) were more stable. This variation is in agreement with the chromosomal locations and dynamics of these three HP1 proteins. HP1γ is associated with active chromatin and mobile, while HP1α and HP1β are typically found in constitutive heterochromatin near (peri) centromeres and are much less mobile (Dialynas et al., 2007). Further indications that heterochromatic loci can display a range of chromatin interaction stabilities dependent upon their precise chromatin composition comes from the observation that stability is modulated by the ratio of HP1α binding and lamin association: interactions for loci that display high levels of HP1α binding but low levels of lamin association are not as stable as those for loci with lower levels of HP1α binding and higher lamin association (Figure S6H).

The differential stability of pair-wise chromatin interactions at different sub-nuclear structures can be directly visualized and quantified by plotting interaction frequencies between pairs of 40 kb loci arranged by their level of factor binding to obtain homotypic interaction saddle plots (Figure 5E). In these plots, pair-wise interactions between loci enriched in factor binding are shown in the lower right corner, and pair-wise interactions between loci not bound by the factor are shown in the upper left corner. For instance, we observe preferential interactions between pairs of loci near speckles, as determined by SON TSA-Seq (Chen et al., 2018). After chromatin fragmentation with DpnII, we observe a progressive loss of preferential interactions between speckle associated loci, while preferential interactions between non-speckle associated loci can be observed even after 16 hours. Conversely, after chromatin fragmentation, we find that preferential interactions between lamin-associated loci remain detectable even at late time points, while preferential interactions between loci not at the lamina disappear relatively fast.

### Chromatin loops dissociate upon chromatin fragmentation

Enriched point-to-point looping interactions are detected as “dots” in Hi-C interaction maps. The majority of these represent interactions between pairs of convergent CTCF sites (Rao et al., 2014). We were interested in determining the fate of such loops after chromatin fragmentation. We aggregated Hi-C data for purified nuclei at pairs of sites that had previously been shown to engage in looping interactions in K562 cells (Rao et al., 2014). We readily detected these loops in intact purified nuclei (Figure 6A). After fragmentation with HindIII for 4 hours, loops remained present and appeared to become slightly stronger. However, fragmenting chromatin with DpnII resulted in loss of loops over time. Although loops appeared somewhat increased at t = 5 minutes after DpnII digestion, they were greatly reduced at t = 1 hour and at later timepoints.

**Figure 6:**
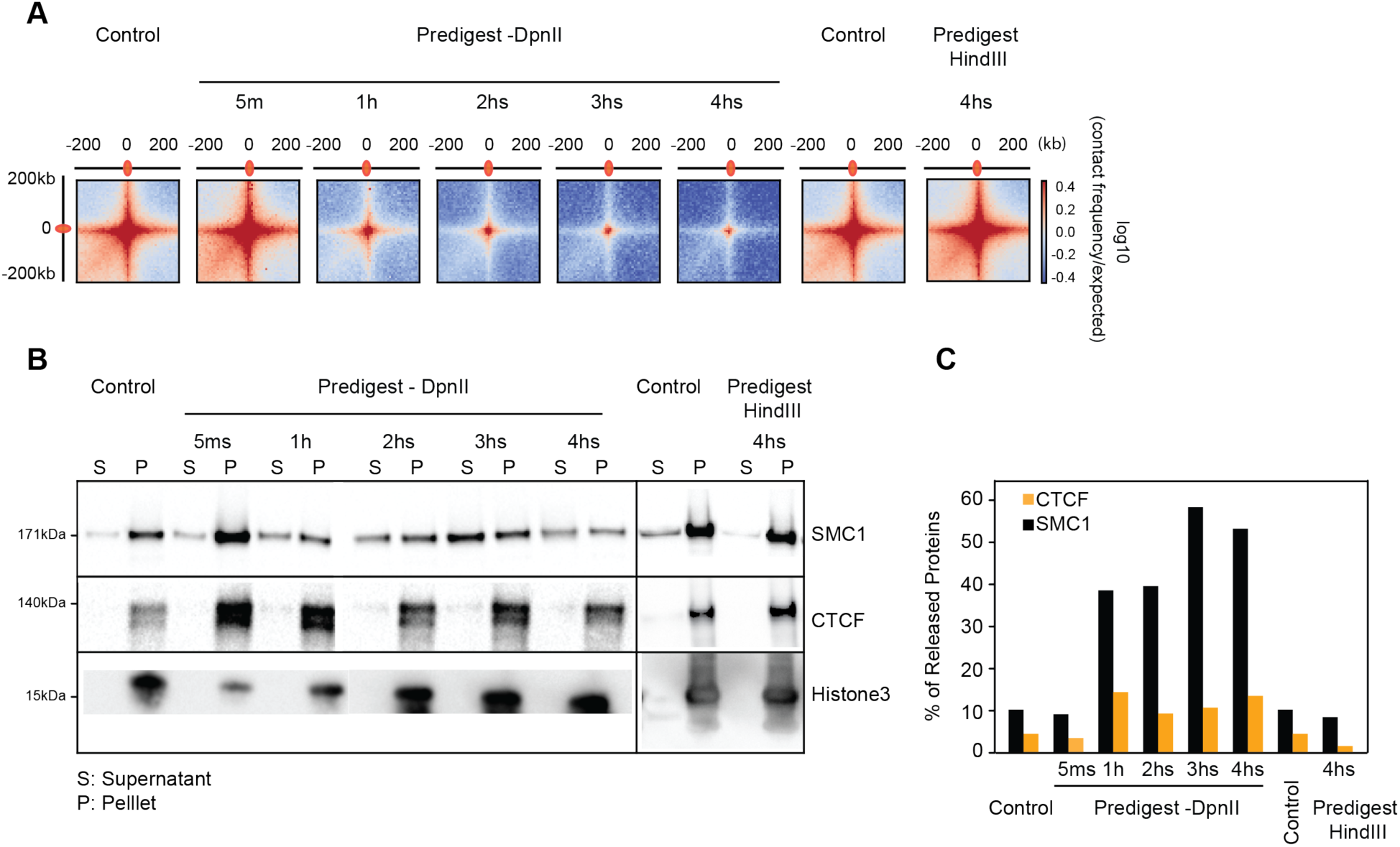
Chromatin loop dissociation upon fragmentation. **(A)** Aggregated distance-normalized Hi-C interactions around 6,057 loops detected in K562 cells by HiCCUPS (Rao et al., 2014) at 10 kb resolution, for control nuclei and nuclei digested with DpnII up to 4 hours, and for nuclei digested with HindIII for 4 hours. **(B)** Western blot analysis of CTCF, cohesin and Histone H3 abundance in soluble and chromatin-bound fractions obtained from control nuclei and from nuclei pre-digested with DpnII up to 4 hours and HindIII for 4 hours. **(C)** Quantification of the data shown in panel B. Percentage of released protein is the ratio of protein level in the soluble fraction divided by the sum of the levels in the soluble and chromatin-bound fractions.

Chromatin loops are thought to be formed by loop extrusion mediated by cohesin that is blocked at convergent CTCF sites. Therefore, we assessed whether CTCF and cohesin binding to chromatin is affected by chromatin fragmentation. We fractionated proteins in chromatin-bound and soluble fractions using the previously described chromatin binding assay ((Liang and Stillman, 1997), Methods). In intact nuclei, most of the CTCF and cohesin is associated with chromatin (Figure 6B, C). Digesting chromatin with HindIII did not lead to dissociation of CTCF or cohesin. However, fragmenting chromatin with DpnII led to dissociation of cohesin after 1 hour, while CTCF binding was only weakly affected. We conclude that DNA fragmentation to <6 kb fragments, but not to 10-25 kb fragments, leads to loss of cohesin binding and loss of looping interactions. These results are consistent with earlier observations that showed that in yeast stable chromatin binding by cohesin requires intact DNA (Ciosk et al., 2000). These data can be interpreted in the context of the model where cohesin rings encircle DNA (pseudo-) topologically (Srinivasan et al., 2018). This model of binding predicts that when DNA is fragmented, the cohesin ring can slide off nearby free ends. Our observation that cohesin binding and loops are disrupted when chromatin is fragmented to <6 kb fragments suggest that loops are maintained by the encirclement of cohesin rings around the loop bases bound by CTCF.

### DISCUSSION

Hi-C interaction maps represent population-averaged folding of chromosomes but do not reveal whether individual pair-wise contacts are dynamic or stable. Liquid chromatin Hi-C reveals chromatin interaction stabilities genome-wide. In liquid chromatin Hi-C chromatin is fragmented prior to fixation. After fragmentation, Hi-C is then used at different time points to determine the extent to which the initially compartmentalized conformation of chromosomes is lost and the formerly spatially separated loci become mixed.

We observe an initial strengthening of A/B compartmentalization following partial digestion. This result supports a “block copolymer” model of chromatin in the interphase nucleus, where B regions of chromosomes tend to cluster together, and A regions cluster together. Partial DNA digestion leads to a strengthening of compartmentalization by removing covalent linkages between A and B blocks, as long as the fragments are still large enough so that attractive forces between them are sufficient for phase segregation.

Further fragmentation of chromatin into pieces that are too short to maintain the phase-separated state leads to progressive dissolution of chromatin conformation. The kinetics of this dissolution process provides insights into the attractive forces between chromatin segments, the intrinsic mobility of loci, the dynamics of nuclear organization, and the protein factors that can mediate chromatin interaction stability. Using liquid chromatin Hi-C we obtain a view of the dynamics of chromatin interactions throughout the nucleus and the genome (Figure 7A). We find that chromatin dissolution is dependent on both chromatin state and fragmentation level. After correcting for fragmentation level we observe that chromatin interactions at different sub-nuclear structures differ in stability, with lamin-associated loci engaged in the most stable interactions and speckle and polycomb associated loci being most dynamic. Further, we find support for the model that CTCF-CTCF loops are stabilized by cohesin rings that encircle loop bases (Figure 7B).

**Figure 7:**
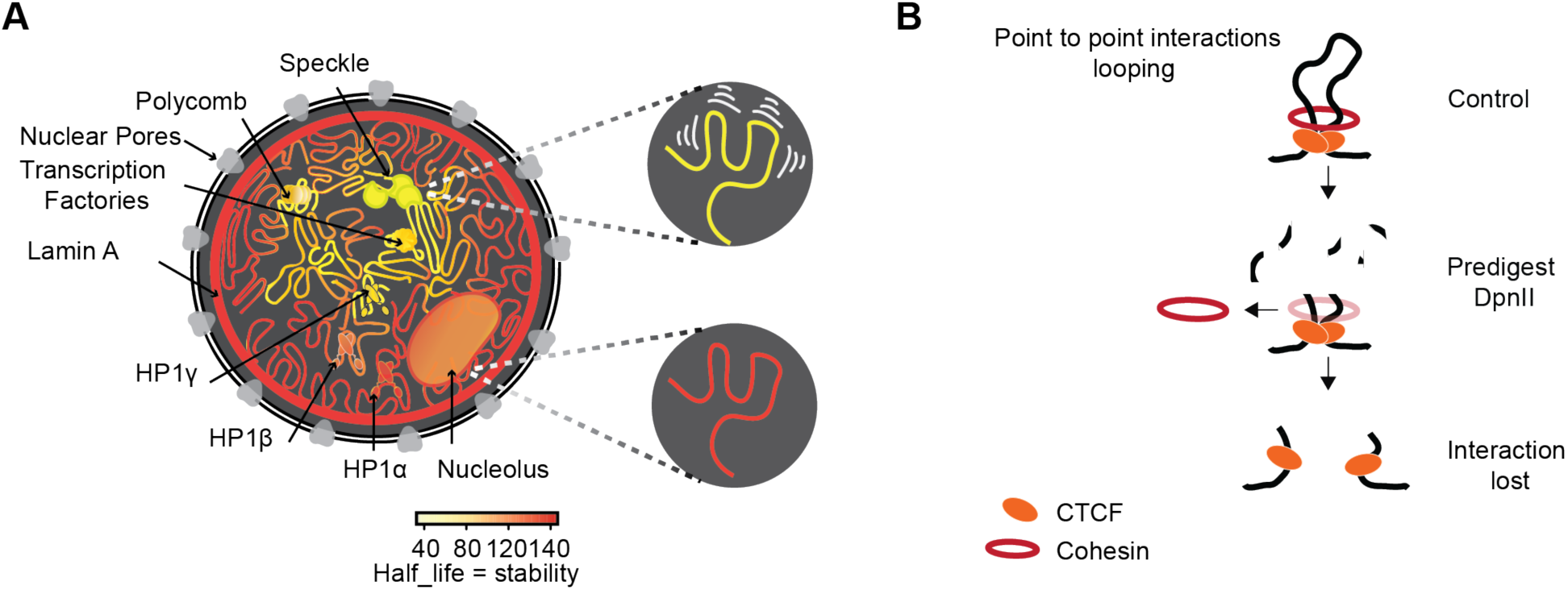
Illustration of chromatin interaction dynamics in the nucleus and model for cohesin loss after chromatin digestion. **(A)** Left: Schematic representation of varying chromatin interactions dynamics at different sub-nuclear domains. Shortest half-life reflects the least stable interactions (yellow), while longest half-life reflects the most stable interactions (dark orange). Nuclear subdomains differ greatly in their stability. Top right: Chromatin anchored at speckles is driven by the most dynamic interactions. Bottom right: Chromatin anchored at the nuclear lamina involves the most stable interactions. **(B)** Model for how cohesin rings stabilize CTCF-CTCF loops by encircling loop bases. Top: Cohesin ring encircles loop bases at convergent CTCF sites. Middle: Pre-digestion with DpnII cuts loop into chromatin fragments <6 kb, and the cohesin ring can slide off nearby ends. Bottom: CTCF remains bound to digested chromatin fragments but interactions between CTCF-bound sites are lost.

Loci that are part of longer intact chromosomes display sub-diffusive dynamics, and their mobility is strongly constrained by the polymeric nature of chromatin and its folded state. In addition, their mobility is modulated by attractive interactions with other loci by factors that themselves are dynamic, and by local chromatin density. Live cell imaging experiments have examined locus motion extensively and found differences in mobility and constrained diffusion dependent on sub-nuclear position and chromatin state and activity (Bronshtein et al., 2009; Bronshtein et al., 2015; Hediger et al., 2002; Marshall et al., 1997; Nagashima et al., 2019; Shinkai et al., 2016; Thakar et al., 2006; Therizols et al., 2010). In such experiments the movement detected is strongly constrained by the fact that loci are part of very long chromosomes. A previous key study, which inspired the current work, aimed to identify the factors that determine intrinsic locus-locus interactions and locus mobility by explicitly removing the strong polymeric constraint due to linkage (Gartenberg et al., 2004). In that work, the mobility of an individual locus and its preference for association with sub-nuclear structures was measured by imaging after excising the locus from its normal genomic location so that it was freed from the polymeric constraint (Gartenberg et al., 2004). Specifically, a silent locus was excised from a yeast chromosome and its intrinsic preference for association with other silent loci and the nuclear periphery was assessed and found to depend on specific protein complexes bound to these loci that are involved in gene silencing. In our liquid chromatin Hi-C experiments, we digest DNA *in situ* and the polymeric constraint on movement is removed for all loci simultaneously, in effect performing a genome-wide variant of the experiments performed by Gartenberg et al. After fragmentation, dynamics of locus mixing is mostly determined by interactions of individual fragments with each other and with sub-nuclear structures mediated by local chromatin-associated factors, histone modifications, and chromatin density.

We show that chromosomal compartmentalization can tolerate genome-wide fragmentation with HindIII in >10-25 kb fragments. Our micro-mechanical elasticity measurements also show that chromosomes remain mechanically fully connected in those conditions. These results indicate that interactions between 10-25 kb fragments are stable enough to maintain the initially phase- or microphase-separated state of the nucleus, at least for several hours. This is in agreement with recent independent locus-specific experiments. First, previous studies had suggested that regions of several hundred kb were long enough to correctly position themselves in vivo according to their chromatin state in the corresponding compartment (van de Werken et al., 2017). Second, super-enhancers that typically range in size between 10 and 25 kb were found to associate both *in cis* and *in trans*, especially in the absence of cohesin (Rao et al., 2017), indicating that interactions between these 10-25 kb elements are stable enough to facilitate their clustering. Third, high-resolution Hi-C analysis in *Drosophila* indicates that domains ∼25 kb in size can phase-separate according to their chromatin status (Rowley et al., 2019; Rowley et al., 2017). Similarly, in *Drosophila*, polycomb-bound loci can cluster together at polycomb bodies when these loci are at least tens of kilobases long (Cheutin and Cavalli, 2012). Combining all these data with our liquid chromatin Hi-C results, we conclude that chromatin phase segregation can occur when domains of a particular chromatin state are at least 10 kb. This number is not far from an independent estimate of the block size required for phase separation based on polymer simulations of preferred interactions between HP1-bound loci (20 kb, (MacPherson et al., 2018)). Notably, digestion studies indicate that mitotic chromosomes are also constrained by stable chromatin interactions spaced by approximately 15 kb (Poirier and Marko, 2002).

Our results obtained with DpnII digestion where the genome is fragmented in <6 kb fragments show that these fragments are too short to maintain stable phase-segregated domains. We find that the stability of interactions between <6 kb fragments, and the rate of mixing of initially segregated loci depends both on the level of DpnII fragmentation and on their chromatin state and association with sub-nuclear structures: interactions at the nuclear lamina are relatively stable, those near nuclear speckles and polycomb complexes are highly unstable, while interactions for loci associated with different heterochromatin proteins and the nucleolus displayed a range of intermediate stabilities. The dynamics of associations between loci are therefore determined by chromatin-associated factors, and may also be determined directly by the biochemical properties of histone tail modifications. For instance, in a recent study, the Rosen lab found that chromatin fragments can form droplets in vitro and that the dynamics of chromatin fragments within these droplets are dependent upon both H1 binding and histone acetylation (Gibson et al., 2019): acetylation of histones resulted in more mobility of fragments and reduced droplet formation, while binding of H1 led to stable droplet with strongly reduced mobility of chromatin fragments. These results are consistent with the dynamics we observe here for active and inactive chromatin fragments within liquid chromatin.

Our results allow a crude estimate of the Flory-Huggins χ parameter for A/B segregation of chromatin in the nucleus. Given that HindIII and DpnII cut chromatin into segments of approximately 15 kb and 6 kb respectively, a reasonable estimate of the minimum length of fragments necessary to drive A/B segregation is *N*\* = 11±4 kb. Given that these fragments are small compared to the A/B compartmentalization scale of a few Mb, the fragments will be essentially A- or B-type (euchromatin or heterochromatin) homopolymers. For homopolymers, the critical length needed for phase separation is *N*\*=2/χ (de Gennes, 1979), indicating χ = 0.18 ± 0.05/kb (χ = 0.03±0.01 /nucleosome). Given that χ is in *k_B_T* units (1 *k_B_T* = 0.6 kcal/mol at room temperature), its small value indicates that the effective demixing interaction between nucleosomes is weak, consistent with a liquid-like phase-separation picture for A and B compartments, where regulation of chromatin organization and compartmentalization is possible by relatively small changes in nucleosome interactions (*e.g.*, via histone modifications). While extremely crude (*e.g.*, we have used the χ estimate for a dense polymer melt rather than for a concentrated solution) our data indicate a weak value for χ, and that the interactions that are driving compartmentalization are a small fraction of a kcal/(mol.nucleosome).

Generally, heterochromatic loci are engaged in the most stable interactions. This implies that these associations play a dominant role in spatial compartmentalization of the nucleus, consistent with predictions made by simulations (Falk et al., 2019). However, there is a range of stabilities correlating with different types of heterochromatin. Interactions between loci at the nuclear laminar are the most stable, suggesting that these loci are firmly tethered to the lamina meshwork. Imaging experiments showed that loci near the lamina can become embedded within the lamina meshwork, and that lamin proteins themselves are also stably localized within the lamina (Broers et al., 1999).

Loci associated with the three different HP1 proteins display different dissociation kinetics. Interestingly, these differences correlate with different dynamics and sub-nuclear locations of the HP1 proteins in the nucleus. HP1γ (CBX3) binds relatively transiently to euchromatin, which may explain the dynamic nature of chromatin interactions between loci bound by this protein (Dialynas et al., 2007). On the other hand, interactions between loci enriched in HP1α (CBX5) are more stable, which likely is the result of more stable HP1α binding to heterochromatin (Strom et al., 2017). Further, the clustering of heterochromatic loci bound by HP1α may be related to condensate formation by HP1α proteins at and around heterochromatic loci (Strom et al., 2017).

Polycomb bound regions of DNA are domains of facultative heterochromatin critical to proper vertebrate and invertebrate development. These regions spatially cluster together to form polycomb bodies visible in cell nuclei (Saurin et al., 1998). Polycomb bodies are shown to compact their associated chromatin as a potential mechanism for gene repression (Boettiger et al., 2016; Francis et al., 2004; Grau et al., 2011). Interestingly, liquid chromatin Hi-C showed differences in chromatin interaction stability between facultative heterochromatic domains marked by polycomb and constitutive heterochromatic domains marked by lamina association or binding of HP1α/HP1β proteins. While many chromatin contacts in constitutive heterochromatin were maintained even after 16 hours of digestion, the half-life for chromatin contacts at polycomb-bound regions was short, on a scale similar to more open and active regions of the genome. Thus, while polycomb domains and constitutive heterochromatin domains can both spatially segregate, when they are long enough, their biophysical nature is very different. The compacted states of polycomb and HP1α bound chromatin appear to form via a similar phase-separation mechanism mediated by multivalent interactions between specific CBX homologs. *In vitro and in vivo,* both CBX2 (polycomb subunit) and CBX5 (HP1α) are capable of forming condensates of polycomb bodies and constitutive heterochromatin, respectively (Larson et al., 2017; Plys et al., 2019; Tatavosian et al., 2019). Our data indicate that these different structures or condensates and associated chromatin have very different properties. Both are dynamic with regards to the exchange of proteins and both have a fraction of stably bound proteins (Strom et al., 2017; Youmans et al., 2018), but the stability of interactions between loci mediated by these factors is distinct, possibly related to differences in affinity between CBX proteins and chromatin: the binding affinity of CBX5 (Hp1α) for its preferential histone modification H3K9me3 is much higher than the affinity of CBX2 for its preferential mark H3K27me3 (Kaustov et al., 2011).

Highly dynamic interactions occurred between loci associated with speckles. This is interesting because previous work had identified speckles to be one of two structural anchors, the other one being the lamina, that determine the organization of the nucleus (Chen et al., 2018; Quinodoz et al., 2018). Our data indicate that these structures differ greatly in how they anchor chromosome conformation: associations at the lamina involve stable interactions while anchoring at speckles is driven by more dynamic interactions.

The dynamics we observe for loci after chromatin fragmentation is likely related to the intrinsic dynamics of those loci while being part of full-length chromosomes. This is confirmed by the fact that active and inactive chromatin are more dynamic and mobile as assessed by both live cell imaging and liquid chromatin Hi-C. Our finding that interactions between active and open chromatin are more dynamic is consistent with observations that treating cells with histone deacetylase inhibitors, which leads to hyperacetylated chromatin, results in a reduced nuclear stiffness as detected by micromechanical measurements (Stephens et al., 2018). The reduced stiffness indicates weaker chromatin – chromatin interactions when chromatin is hyperacetylated.

It is important to point out that during the liquid chromatin Hi-C procedure, some chromatin factors may dissociate from the nucleus, and this could affect the locus mixing behavior we observe. This will likely be the case for proteins that rapidly dissociate. As we discussed above, protein binding dynamics, e.g., of HP1 proteins, may determine the stability of chromatin interactions. An example of proteins that dissociate from chromatin upon fragmentation is the cohesin complex, possibly due to the way it binds chromatin by encircling chromatin fiber(s). However, the cohesin complex is not involved in compartment formation (Haarhuis et al., 2017; Nora et al., 2017; Nuebler et al., 2018; Rao et al., 2017; Schwarzer et al., 2017; Wutz et al., 2017), and thus its dissociation may not influence the stability of phase separation-driven chromatin interactions detected by liquid chromatin Hi-C.

The current work analyzed the intrinsic chromatin interaction strengths and dissolution kinetics of chromosome conformation within otherwise inactive nuclei and how these measurements along chromosomes relate to chromatin state and chromosomal compartmentalization. Future work can focus on how these kinetic properties change in cells or nuclei, where active processes such as transcription, replication, chromatin compaction and condensation, and loop extrusion are also acting, and on determining the roles of RNAs, protein complexes, and histone modifications in modulating the attractive forces between loci and the dynamics of genome folding in general.

## Acknowledgements

J.D. and J.F.M. acknowledge support from the National Institutes of Health Common Fund 4D Nucleome Program (U54-DK107980). This work was also supported by a grant from the National Human Genome Research Institute (NHGRI) to J.D. (HG003143), another NHGRI grant to Z.W. (HG009446), and by grants from the National Cancer Institute (U54-CA193419) and from the National Institutes of Health (R01-GM105847 to J.F.M. and K99-GM123195 to A.D.S.). J.D. is an investigator of the Howard Hughes Medical Institute. We thank Leonid A. Mirny and Edward J. Banigan for discussions and Jian Ma for sharing K562 sub-compartment assignments.

## Author contributions

J.D conceived the project. H.B. performed all 3C, 5C, Hi-C and liquid chromatin Hi-C and chromatin fractionation experiments. DLL performed restriction digestion efficiency (DpnII-seq) experiments. A.D.S. performed micromechanical studies and analyzed the data. T.B. and H.B. analyzed data. S.V. contributed analysis tools for liquid chromatin Hi-C analysis. J.F.M. provided polymer scaling ideas relevant to data interpretation. All authors contributed to writing the manuscript.

## Data availability

All data will be made publicly available upon publication via GEO and the 4D Nucleome Data Coordination Center. <u>Submission of the all sequencing data to GEO is in progress and the accession number will be made available to the reviewers as soon as we have the number and secure accession key.</u>

## Code availability

All code for data processing and analysis, described in detail in the Methods, is available through the following GitHub accounts:

https://github.com/dekkerlab/5C-CBFb-SMMHC-Inhib

https://github.com/dekkerlab/cMapping

https://github.com/dekkerlab/cworld-dekker

https://github.com/tborrman/DpnII-seq

https://github.com/tborrman/digest-Hi-C

https://github.com/hms-dbmi/hic-data-analysis-bootcamp

https://github.com/mirnylab/cooltools/tree/master/cooltools

## METHODS

### Digestion, cross-linking and copolymer architecture and hetero/euchromatin phase separation

We have found that moderate digestion leads to formation of stronger inter-compartment interactions in Hi-C. While this may seem somewhat paradoxical at first glance, this effect is rather straightforward to explain if we remember two basic physical facts about chromatin in the G1 nucleus. First, individual chromosomes have a “blocky” structure, with hundreds of kilobase-scale stretches of alternatingly heterochromatin-like and euchromatin-like character along their length. Second, the chromatin is in a state of quite extensive “crosslinking” (i.e., noncovalent chromatin-chromatin interactions mediated by proteins including gene regulatory factors such as cohesin, and heterochromatin linkers such as HP1α.

Therefore, chromatin in the G1 nucleus can be considered as a set of blocks of euchromatin and heterochromatin (the A and B compartments consisting of regions of predominantly euchromatin vs heterchromatin, respectively), which are constrained to be near each other by being part of the same linear chromosomes, i.e., effectively being long many-block copolymers. We suppose that the A and B heterochromatin/ euchromatin monomers have a weak tendency to repel one another (or equivalently that A-A or B-B attract one another, for example via protein-mediated nucleosome-nucleosome interactions acting preferentially on euchromatin or heterochromatin, or even via physio-chemical effects such as relative hydrophobicity of more methylated nucleosomes).

If we suppose the A and B blocks to be on average N monomers long (roughly nucleosomes for the sake of this discussion), then under melt-like conditions the standard Flory theory of polymer phase separation predicts that if we were to cut the polymers into pure A and B blocks at the block boundaries (i.e., at a spacing of N monomers commensurate with the block sizes), they would phase separate for a segment-segment interaction strength stronger than chi* = 2/N (de Gennes, 1979). Note that this level of interaction (given approximately in k_B T units) is proportional to 1/N where N is crudely in nucleosome units; for 200 kilobase blocks, we have approximately N=1000, indicating that small fractions of a k_B T in effective A/B repulsion or A-A or B-B attraction is sufficient to drive strong euchromatin/heterochromatin phase separation (Marko and Siggia, 1997).

Now if we were to instead cut less frequently than this, say at every second block boundary, so as to arrive at a system of AB linear diblock copolymers each of length 2N (N monomers of A followed by N monomers of B), the constraint that the A and B blocks be connected suppresses phase separation, increasing the critical interaction (all other factors held constant) to chi* = 5.3/N (Leibler, 1980). In this case bulk phase separation cannot occur, but instead local, or “microphase separation” occurs, with formation of micelle-like or layered phase-separated structures. Nevertheless, for chi >> chi*, strong segregation of the A and B monomers can still occur.

If we were to not cut at all, but rather to suppose that the chromosomes are very long multiblock copolymers, with many blocks each of N monomers alternating between A and B (“ABABABAB… multiblock copolymers”), the critical interaction strength will rise with increasing number of blocks, approaching the limit chi* = 7.5/N for many blocks (Matsen and Schick, 1994). Therefore, starting from this limit, the tendency for chromosome domains to phase separate will be enhanced by cutting the chromosomes up into successively smaller pieces: as chromatin cutting increases from no cutting, we expect to see intensification of A/B compartment contrast in the Hi-C map.

Now, if we cut too frequently, when the cuts become spaced smaller than the block size (cut spacing M < N monomers), we will have the situation that the critical interaction strength will become chi* = 2/M > 2/N, i.e., the cuts are frequent enough to suppress phase separation by decreasing the amount of interaction enthalpy per polymer “molecule”. Therefore we expect that overly frequent cutting will cause a reduction in A/B compartment Hi-C map contrast, i.e., for some intermediate level of cutting similar to the sizes of the A and B blocks, one will see a maximum level of A/B compartment contrast.

There is also likely an effect of “crosslinking” (“chromatin cross-bridging”), which provides an additional level of constraint suppressing phase separation, above the linear-multiblock architecture of chromosomes. For example, taking linear diblock copolymers (N A monomers followed by N B monomers) and circularizating them raises the critical interaction for microphase separation from 5.3/N to 8.9/N, nearly a factor of 2 (Marko, 1993).

Similarly, if we start with A and B homopolymers each of length N, constraining them to have their ends at a flat surface, thus forcing them to mix at the surface, increases the critical interaction for phase separation from 2/N up to 4.5/N (Marko and Witten, 1991), with microphase separation again occurring in the constrained case. Releasing chromatin crosslinking/cross-bridging constraints (which also will occur for chromatin cutting) will in general also reduce the interaction strength needed to drive phase separation, increasing A/B compartment contrast in Hi-C maps.

In conclusion, basic polymer phase separation theory predicts that gradually increasing the cleavage of chromatin will gradually increase the intensity of A/B compartment contrast in Hi-C maps until the cuts are spaced by approximately one A or B block; further cutting will reduce the intensity of phase separation and A/B compartment contrast. Notably, the nature of the segregation can be expected to be “microphase segregation” rather than bulk phase separation, until the number of cuts is sufficient to liberate A or B “homopolymer” segments.

### K562 nuclei purification

Three sucrose cushions were made before starting nuclei purification. 30 mL of 30% sucrose [10 mM PIPES pH 7.4, 10 mM KCl, 2 mM MgCl2, pH adjusted to 7.4 using 1 N KOH, 30% sucrose, 1 mM DTT (added prior to use), 1:100 protease inhibitor (Thermo Fisher 78438) (added prior to use)] was transferred to a 50 mL tube, then 5 mL of 10% sucrose [10 mM PIPES pH 7.4, 10 mM KCl, 2 mM MgCl2, 10% Sucrose, 1 mM DTT (added prior to use), 1:100 protease inhibitor (added prior to use)] was slowly loaded in top of 30% sucrose, and the tubes were incubated at 4°C until needed. K562 cell pellets (100 million cells) were lysed using the following nuclear isolation procedure. After the cells were spun, the pellets were washed twice with 10 mL HBSS, then pelleted after each wash at 300 rpm for 10 min at 4°C. Cell pellets were dissolved in 15 mL nuclear isolation buffer [10 mM PIPES PH 7.4, 10 mM KCl, 2 mM MgCl2, 1 mM DTT (added prior to use), 1:100 protease inhibitor (added prior to use)], pH adjusted to 7.4 using 1 M KOH]. Then, cells were lysed on ice in a 15 mL Dounce homogenizer with pestle A (KIMBLE Kontes 885002-0015) by moving the pestle slowly up and down 20 times, followed by incubation on ice for 20 min and another 20 strokes. Next, each 5 mL of lysed extract was loaded slowly on top of a sucrose cushion prepared earlier. Then the tubes were spun for 15 min at 800 g at 4°C. The supernatant was removed carefully for a good recovery of the nuclei pellet in the bottom of the tube. Nuclei pellets were resuspended in 1 mL of HBSS, then spun for 5 min at 5,000 g at 4°C using a benchtop refrigerated centrifuge. Then, the nuclei pellet was resuspended in 3 mL HBSS, and 1 µL was taken to quantify the nuclei before the 3 mL was split over two microfuge tubes and spun for 5 min at 5,000 g at 4°C using a benchtop refrigerated centrifuge. Finally, the nuclei pellet was dissolved into an adequate total volume to obtain 1 million nuclei per 0.1 mL of Nuclei storage buffer (NSB) [10 mM PIPES pH 7.4, 10 mM KCl, 2 mM MgCl2, 50% glycerol, 8.5% sucrose, 1 mM DTT (added prior to use), 1:100 protease inhibitor (added prior to use)]. Each 0.5 mL of NSB containing 5 million nuclei was transferred to a microfuge tube and stored at −80°C.

### 3C (Chromosome Conformation Capture)

3C was performed as described in “From cells to chromatin: Capturing snapshots of genome organization with 5C technology” (Ferraiuolo et al., 2012).

**Crosslinking**: 1.25 mL of 37% formaldehyde was added to 40 mL of HBSS. 50 million cells or nuclei were washed twice using 20 mL of HBSS and then pelleted at 500 g for 10 min. The pellet was resuspended in 5 mL HBSS and then added to 41.25 mL of HBSS and formaldehyde (final formaldehyde concentration was 1%). The sample was incubated at RT for 10 min on a rocking platform. Afterward, to stop cross-linking 2.5 mL of 2.5 M glycine was added and samples were incubated at RT for 5 min on a rotating platform. To pellet the crosslinked cells or nuclei the sample was centrifuged at 800 g for 10 min at 4°C. After discarding the supernatant the pellet was washed twice using HBSS. Next, the pellet was either processed immediately as described below or was stored at −80 °C after flash freezing using liquid nitrogen.

**Cell lysis**: (This step was included when cells are used, but was skipped for 3C with purified nuclei).Cells were lysed by adding 2 mL of cold lysis buffer [10 mM Tris-HCl (pH=8.0), 10 mM NaCl, 0.2% Igepal CA-630 (NP40)] and 20 µL of 100x Protease inhibitors. The sample was incubated on ice for 15 min to let the cells swell. The cells were lysed on ice using the homogenizer with pestle A (KIMBLE Kontes 885300-0002) by moving the pestle slowly up and down 30 times and incubating on ice for 1 min followed by another 30 strokes. The sample was transferred to two 1.5 mL microcentrifuge tubes, spun at 5,000 g at RT for 5 min using a benchtop centrifuge.

**Digestion**: each pellet was washed using 1 mL cold 1X NEBuffer 2.1, then spun at 5,000 g for 5 min at RT using a benchtop centrifuge, afterward each pellet was resuspended in 250 µL of 1X NEB2.1 buffer, and the two pellets were pooled (∼ 500 µL). 50 µL aliquots of the suspension were transferred to 10 new 1.5 mL microfuge tube and 292 µL of 1x NEBuffer 2.1 was added to each tube. Next, 38 µL of 1% SDS was added per tube and mixed well, the samples were incubated at 65°C for 10 min, then placed on ice. 44 µL of 10% Triton X-100 was added to each tube to quench SDS. Finally, 400 U of EcoRI (NEB R0101L) was added per tube and incubated at 37°C overnight on a thermocycler (with 900 rpm for 30 sec every 4 min).

**Ligation:** 86 µL of 10% SDS was added to the digested samples and the samples were then incubated at 65°C for 30 min for EcoRI inactivation after which the tubes were placed on ice. Each sample was then transferred to a 15 mL conical tube and 7.69 mL of ligation mix was added [820 µL 10% Triton X-100, 820 µL 10x ligation buffer (500 mM Tris-HCl pH7.5, 100 mM MgCl2, 100mM DTT), 82 µL 10 mg/mL BSA, 82 µL 100 mM ATP and 5.886 µL ultrapure distilled water]. Finally, 10 U of T4 ligase (Invitrogen 15224090) was added per tube before incubation at 16°C for 2 hr on a thermocycler (with 900 rpm for 30 sec every 4 min).

**Reverse Crosslinking**: 50 µL of 10 mg/mL proteinase K (Fisher BP1750I-400) was added per tube, the sample was incubated at 65°C for 4 hr followed by a second addition of 50 µL 10 mg/mL Proteinase K and overnight incubation at 65°C on a thermocycler (with 900 rpm for 30 sec every 4 min).

**DNA purification:** Tubes were cooled at room temperature, at this stage each tube contains ∼ 8.21 mL final volume. The samples from every two tubes were combined to a 50 mL conical tube (∼16,42 mL) to have five tubes in total. DNA was extracted by adding an equal volume of 17 mL of saturated phenol pH8.0: chloroform (1:1) (Fisher BP1750I-400) and vortexing for 3 min. Then the mix was transferred to a 15 mL phase-lock tube (Quiagen 129073) followed by spinning tubes at 5,000 g for 10 min. The upper phase was taken to a 50 mL tube to start the second extraction. We added an equal volume of 17 mL saturated phenol pH 8.0: chloroform (1:1), vortexing for 1 min. Then the upper phase was transferred to a 15 mL phase-lock tube, and tubes were centrifuged at 5,000 g for 10 min. We pooled all the upper phases from all 5 tubes ∼ 85 mL into a single 300 mL high-speed centrifuge tube to precipitate the DNA. 8.5 mL (1/10 volume) of 3M sodium acetate pH 5.2 was added and brief vortexing was performed, then 212 mL (2.5 volumes) of ice-cold 100% ethanol was added, and the tube was inverted slowly several times and incubated at −80° C for 1 hr. Afterward, the DNA was pelleted at 16,000 g for 30 min at 4°C. The supernatant was discarded and the pellet was dissolved in 500 µL 1X TLE and transferred to a 0.5 mL AMICON Ultra Centrifuge filter (UFC5030BK EMD Millipore). The column was centrifuged for 5 min at 14,000 g and the flow-through was discarded. The column was washed 4 times using 450 µL of 1X TLE for desalting DNA. After the final wash, the library remaining in the column (∼50 µL) was eluted in 30 µL of 1XTLE, the column was flipped upside down into a new tube to collect DNA by centrifugation for 3 min at 4,000 g. RNA was degraded by adding 1 µL of 10 mg/mL RNAase A and incubation for 30 min at 37°C.

**Quality control assessment**: to test the quality of the 3C library we used PCR to amplify a specific ligation product formed by two nearby restriction fragments, using the following primers: GPF33: GACCTCTGCACTAGGAATGGAAGGTTAGCC GPF23: GACTAATTCCTGACACTACTTGAGGGATAC

The amplicon was digested with EcoRI to assess the efficiency of 3C ligation. 3C primers used for analysis of the beta-globin locus are listed in Supplemental Table S1.

### BAC library for 3C-PCR

BAC DNA was generated as described (Dostie et al., 2006). A control ligation library covering the Beta-globin locus (ENCODE region ENm009) was generated using BACs overlapping the region. Starting with a mixture of DNA of seven BACs (CTC-775N13, RP11-715G8, CTD-3048C22, CTD3055E11, CTD-2643I7, CTD-3234J1, and RP11-589G14) (Invitrogen), mixed in equimolar ratios, we used the same steps described in the 3C protocol above starting from the digestion step. BAC clones were digested with EcoRI, then randomly ligated, and the DNA was purified. The BAC ligation library reflects random ligation of EcoRI fragments throughout the beta-globin locus, so any difference in PCR signal for 3C primer pairs along the beta-globin locus due to differences in primer efficiency can be corrected by normalizing the amount of PCR product obtained with the 3C library to the amount obtained with the BAC ligation library.

### Chromosome Conformation Capture Carbon Copy (5C)

#### Experimental design

Probes were designed as described (Dostie et al., 2006). 213 5C probes were designed for a ∼1 Mb region (chr11:4730996 −5729937; hg18) around the Beta-globin locus at EcoRI restriction sites using publicly available 5C primer design tools (Lajoie et al., 2009). Probes were designed according to a single alternating scheme exactly as described before (Lajoie et al., 2009) and the genomic uniqueness of all primers was verified with the SSAHA algorithm. For each EcoRI fragment at the 1 Mb target region a primer was designed. 104 5’ forward (FOR) and 109 5’ reverse (REV) primers were designed. 5C primers are listed in Supplemental Table S2.

#### Generation of 5C libraries

5C libraries were generated as described before (Ferraiuolo et al., 2012), with three modifications. First, we skipped the gel purification after the adaptor ligation and replaced this with a 1:1 Ampure step to remove unligated DNA and adaptors. Second, barcoded Illumina adaptors were used. Third, we performed the final PCR using TruSeq DNA LT kit Set A (REF 15041757).

**Annealing**: The 5C probes were pooled and combined with the 3C template each reaction contained 800,000 genome copies of 3C template and 0.2 fmol per 5C probe [800,000 genome copies of 3C template, 2 µL of 10X NEB4 (NEB B7004S), 2.75 µL of Salmon Sperm DNA (250 ng; (Invitrogen™ 15632011), 0.25 µL of 1 fmol/µL probes, up to 20 µL ultrapure distilled water]. We set up 8 annealing reactions for each library in a 96-well PCR plate. We then incubated the samples in a PCR machine and ran the following program [95°C for 9 min, Ramp 0.1°C/sec to 55 C, then keep at 55°C for 12 hr].

**Ligation**: We ligated 5C probe pairs, which represent a specific ligation junction in the 3C library, by adding 20 µL of ligation mix [2 µL of [10X *Taq* DNA ligase buffer (NEB B0208S), 0.25 µL Taq DNA ligase (NEB M0208S), 17.75 uL ultrapure distiller water] while the samples are kept in the PCR block at 55°C. We then incubated the reactions for 1 hr at 55°C followed by a 10 min incubation at 65°C; samples were then cooled to 4°C. Negative controls (no ligase, no template, no 5C oligonucleotide) were included to ensure the absence of any contamination.

**PCR amplification**: Universal emulsion primers were used for amplification of the ligated product by using 5C forward and reverse emulsion primers [Forward_primer: CCTCTCTATGGGCAGTCGGTGAT. Reverse_primer : CTGCCCCGGGTTCCTCATTCTCT] for 25 PCR cycles [6 µL of ligation product, 2.5 µL of 10XPCR (600 mM Tris-SO_4_, pH 8.9, 180 mM (NH_4_)_2_SO_4_), 1.8 mM MgCl2, 0.2 mM dNTP, 0.5 µL F-emulsion primer (80 µM), 0.5 µL R-emulsion primer (80 µM), 0.225 µL *AmpliTaq*Ò *Gold* DNA polymerase, ultrapure distilled water to bring volume up to 25 µL]. We then amplified DNA using this PCR program: [95° 9 min, 25 cycles (95°C 30 s, 65°C 30 s, 72° 30 s), 95°C 30 s, 65°C 30 s, 72°C 8 min, 4°C].

We pooled all the PCR reactions for the same library together and concentrated the DNA to 50 µL using 0.5 mL AMICON Ultra Centrifuge filter (UFC5030BK EMD Millipore). DNA was then loaded on a 2% agarose gel, along with a low molecular weight ladder, and the gel was run in a 4°C room at 200 volts for 90 min. The 150 bp DNA that corresponded to the ligated 5C probes was isolated from the gel using the QIAquick Gel Extraction Kit Protocol (QIAGEN 28115). DNA was finally eluted in 32 µL of 1XTLE.

**A-tailing**: A dATP was added to the 3’ ends of the 5C library by adding 18 µL of A-tailing mix [5 µL NEB buffer 2.1, 10 µl of 1 mM dATP, 3 µL Klenow exo (NEB M0212S)] to the 32 µL of DNA sample from the previous step. The reaction was then incubated in a PCR machine [at 37°C for 30 min, then at 65°C for 20 min, and finally cooled down to 4°C]. Next, the tube was placed on ice immediately. 1:1 Ampure was used to remove unligated adaptors. The DNA was finally eluted in 40 µL 1X T4 DNA Ligase buffer (Invitrogen).

**Illumina adapter ligation and paired-end PCR**: For this step, we used the TruSeq DNA LT kit Set A (REF 15041757). 10 µL of ligation mix [5 µL Illumina paired-end adapters, 3 µL T4 DNA ligase Invitrogen, 2 µL 5x T4 DNA ligase buffer (Invitrogen 5X)] was added to the 40 µL sample from the previous step. The ligation sample was then incubated at RT for 2 hours on a tube rotator. Afterward, the sample was run on a 2% agarose gel in a cold room 4°C at 150 volts for 120 min along with a low molecular weight ladder. The 270 bp band that corresponds to 5C products (150 bp) ligated to the two adaptors (64 bp) was extracted from the gel and isolated using the QIAquick Gel Extraction Kit (QIAGEN 28115). DNA was finally eluted in 30 µL 1XTLE.

#### Pre-digestion of nuclei (liquify chromatin)

Purified nuclei as described above (**K562 nuclei purification**) were placed on ice and 1 mL of HBSS was added to each 0.5 mL of 5 million frozen nuclei. After thawing, nuclei were centrifuged 5 min at 5,000 g. The nuclei pellet was washed twice with 1XNEB3.1 for nuclei that would be digested with DpnII or 1XNEB2.1 for nuclei that would be digested with HindIII. The nuclei were pelleted for 5 min at 5,000 g after each wash.

**Isolated nuclei**: a sample of 5 million nuclei was resuspended in 1,250 µL of 1X NEB3.1 as control, and then processed immediately for Hi-C starting at the crosslinking step (see below Hi-C 2.0 protocol).

**Undigested nuclei**: Each sample of two million nuclei was resuspended in 500 µL of 1X NEB3.1 on ice, as and control for the pre-digestion and then treated as described immediately below.

**DpnII pre-digestion**: Each sample of two million nuclei was resuspended in 500 µL of 1X NEB3.1 on ice. Next, 120 U of DpnII (NEB R0543S) was added to the sample in order to obtain 10 U DpnII/µg DNA and then treated as described immediately below.

**HindIII pre-digestion:** Each sample of two million nuclei was resuspended in 500 µL of 1X NEB2.1 on ice. Next, 600 U of HindIII (NEB R0104T) was added to the sample in order to obtain 50 U HindIII/µg of DNA and then treated as described immediately below.

Next, control and pre-digestion samples were incubated at 37°C on a thermocycler (900 rpm for 30 sec every 4 min) for 5 min up to 16 h. Afterward, samples were placed on ice for 10 min.For DpnII-seq and assessment of fragmentation level, a final volume of 10 mM of EDTA was added to inactivate the endonuclease, followed immediately by the DpnII-seq protocol (details of protocols below. DpnII-Seq) or DNA purification for fragment analyzer analysis. For Hi-C, we proceeded immediately to the first step of the protocol (crosslinking as described below). For microscopy, nuclei samples were cross-linked with a 4% final concentration of paraformaldehyde.

### Hi-C 2.0

Hi-C was performed as described (Belaghzal et al., 2017) with some modifications in the crosslinking and lysis step as described below.

**Crosslinking**: isolated, undigested, and pre-digested (with liquified chromatin) nuclei were not pelleted after the pre-digestion step above but were crosslinked immediately as follows: for each sample 1,250 µL volume of nuclei in the digestion buffer was transferred to a 21.875 mL mix [625 µL of 37% formaldehyde + 21.25 mL of HBSS]. For intact cells: 5 million K562 cells or nuclei were washed twice with 15 mL of HBSS and pelleted at 300 g for 10 min, then resuspended in 2.5 mL of HBSS. The sample was transferred to 20.625 mL crosslinking mix [625 µL of 37% formaldehyde + 20 mL of HBSS].

All samples were incubated at RT for 10 min on a rocking platform. Next, to stop cross-linking 1.25 mL of 2.5 M glycine was added to each sample and the mix was incubated at RT for 5 min on a rocking platform. To pellet the crosslinked cells/nuclei, the sample was centrifuged at 1,000 g for 10 min at 4°C. The supernatant was discarded and the pellet was washed twice with HBSS before going to the next step or storing samples at −80°C.

**Cells lysis**: This step is not needed for isolated, undigested, and pre-digested (with liquified chromatin) nuclei. For Hi-C with intact cells: the 5 million crosslinked cells were lysed by adding 1 mL cold lysis buffer [10 mM Tris-HCl (pH=8.0), 10 mM NaCl, 0.2% Igepal CA-630 (NP40)] and 10 µL of 100X Protease inhibitors. The sample was incubated on ice for 15 min to let the cells swell. The cells were lysed on ice using a dounce homogenizer with pestle A (KIMBLE Kontes 885300-0002) by moving the pestle slowly up and down 30 times and incubating on ice for 1 min followed by another 30 strokes. The sample was transferred to a new 1.5 mL microcentrifuge tube, and the sample was centrifuged at 5,000 g at RT for 5 min.

**Digestion**: from each sample (isolated undigested, and pre-digested (with liquified chromatin) nuclei and lysed cells) the pellet was resuspended in 500 µL of ice-cold 1X NEBuffer 3.1, and pelleted for 5 min at 4,000 g. The pellet was washed twice using 500 µL of ice-cold 1X NEBuffer 3.1. After the last wash, the pellet was resuspended in 350 µL 1X NEBuffer 3.1, and 8 µL was taken and kept at 4°C to assess the DNA integrity later. 38 µL of 1% SDS was added to 342 µL (380 µL total volume), and the mixture was resuspended and incubated for 10 min at 65°C. The tube was placed on ice immediately afterward. Next 43 µL of 10% Triton X-100 was added and the sample was mixed gently by pipetting. The tubes were placed at room temperature and 12 µL of 10X NEBuffer 3.1 was added. Then 400 U of DpnII (R0543L) was added and mixed gently before an overnight incubation at 37°C on a thermocycler (with 900 rpm for 30 sec every 4 min). **Biotin Fill-in**: After overnight digestion, the sample was incubated at 65°C for 20 min in order to inactivate the restriction enzyme. Then, 10 µL of the digested sample was taken and kept at 4°C to assess the digestion efficiency later. DNA ends were marked with biotin-14-dATP by adding 60 µL of biotin fill-in master mix [1XNEB 3.1, 0.25 mM dCTP, 0.25 mM dGTP, 0.25 mM dTTP, 0.25 mM biotin-dATP (ThermoFisher#19524016), 50U Klenow polymerase Polymerase I (NEB M0210L)]. Next, the sample was incubated for 4 h at 23°C on a thermocycler (with 900 rpm for 30 sec every 4 min). Finally, the sample was placed on ice immediately for 15 min before proceeding to the next step.

**Ligation:** After fill-in, the total sample volume was ∼535 µL. Ligation was performed by adding 665 µL of ligation mix [240 µL of 5x ligation buffer (1.8X) (Invitrogen), 120 µL 10% Triton X-100, 12 µL of 10 mg/mL BSA, 50 µL T4 DNA ligase (Invitrogen 15224090), and 243 µL ultrapure distilled water (Invitrogen)], to make a total volume of 1,200 µL. The reaction was then incubated at 16°C for 4 hours in a Thermomixer with interval shake.

**Reverse Crosslinking**: 50 µL of 10 mg/mL **proteinase K (Fisher BP1750I-400) was added after ligation, the sample was incubated at 65°C for 4** hr followed by a second addition of 50 µL 10 mg/mL Proteinase K and overnight incubation 65°C

**DNA purification:** Reactions were cooled to room temperature and the 1.3 mL total volume was transferred to a 15 mL tube. The DNA was extracted by adding an equal volume of 1.3 mL of saturated phenol pH 8.0: chloroform (1:1) (Fisher BP1750I-400) and vortexing for 1 min. Then the total volume of 2.6 mL was transferred to a 15 ml phase-lock tube (Quiagen #129065) and tubes were centrifuged at 5,000 g for 10 min. The upper phase was transferred to a 15 mL tube to start the second extraction. An equal volume of 1.3 mL saturated phenol pH8.0: chloroform (1:1) was added and the sample was vortexed for 1 min. Then the mix was transferred to 15 ml phase-lock tube (Quiagen #129065) followed by spinning tubes at 5,000 g for 10 min. The upper phase of ∼1.3 mL was transferred to a 15 mL tube (high speed) to precipitate the DNA. 1/10 volume (130 µL) of 3 M sodium acetate pH 5.2 was added and the sample was briefly vortexed. Then, 2.5 volumes of ice-cold 100% ethanol 3.25 mL was added, the tube was inverted slowly several times and then incubated at −80° C for 1hr. Next, the DNA was pelleted at 16,000 g for 30 min at 4°C. The supernatant was discarded and the pellet was dissolved in 500 µL 1X TLE and transferred to a 0.5 mL AMICON Ultra Centrifuge filter (UFC5030BK EMD Millipore). The column was spun for 5 min at 14,000 g and the flow-through was discarded. The column was washed 4 times using 450 µL of 1X TLE for desalting of DNA. After the final wash the DNA remaining in the column (∼50 µL) was eluted in 52 µL of 1XTLE. The column flipped upside down into a new tube to collect DNA and spun for 3 min at 4,000 g, the volume was adjusted to 102 µL. RNA was degraded by adding 1 µL of 10 mg/mL RNAase A and incubation for 30 min at 37°C. To quantify the DNA concentration, 2 µL of the final DNA sample along with the first 8 µL sample taken before digestion, the 10 µL sample taken after digestion, and various amounts of the 1 kb ladder (NEB#N3232s) were run on 1% Agarose gel.

**Removal of Biotin from unligated ends:** To remove biotinylated nucleotides at DNA ends that did not ligate, the Hi-C sample was treated with T4 DNA polymerase. For each Hi-C sample, we assembled the following reaction: [up to 5 µg of Hi-C library, 5 µL 10x NEBuffer 3.1, 0.025 mM dATP, 0.025 mM dGTP and 15 U T4 DNA polymerase (NEB # M0203L). The samples were brought up to 50 µL total volume adding ultrapure distilled water. Reactions were incubated at 20°C for 4 hours, the enzyme was then inactivated by incubation of the reaction for 20 mins at 75°C and placed at 4°C. Next, the samples were pooled and the volume was brought up to 130 µL 1XTLE in preparation for sonication.

**Sonication:** the DNA was sheared to a size of 100-300 bp using a Covaris instrument [Duty cycle 10%, Intensity 5, Cycles per Burst 200, set Mode Frequency sweeping, continuous degassing, process time 60 sec, Number of cycles] for 3 cycles. The volume was brought up to 500 µL using TLE for Ampure fractionation.

**Size fractionation using AMpure XP:** 500 µL AMpure beads (Beckman Coulter A63881) were added to a 1.5 mL tube labeled as 1.1X. Then the tube was placed on the Magnetic Particle Separator (MPS) for 5 min, and the supernatant was removed. Beads were resuspended in 150 µL AMpure mixture in order to make the 1.1X solution. 400 µL of AMpure mixture was added to 500 µL of sonicated DNA from the previous step and the tube was labeled 0.8X. The sample was vortexed and spun down briefly followed by incubation at RT for 10 min on a rotating platform. Then the tube was placed on the MPS for 5 min at RT. The supernatants were collected and added to the 1.1X tube, the tube was briefly vortexed and spun down followed by incubation at RT for 10 min on a rotating platform. Then the tube was placed on the MPS for 5 min at RT. The supernatant was discarded and the beads in 0.8X and 1.1X tubes were washed twice with 1 mL 70% ethanol. Beads were reclaimed by the MPS for 5 min. Beads were then air-dried on the MPS until ethanol had evaporated completely. Next, 51 µL of 1XTLE was added to the 0.8X and 1.1X tubes to resuspend the DNA from the beads. Tubes were incubated at RT on a rotating platform for 10 min. Then the tubes with AMpure beads from both 0.8X and 1.1X tubes were placed on the MPS for 5 min. Finally, the supernatants were transferred to 1.7 mL tubes labeled 0.8X and 1.1X. Our sample with DNA that ranges from 100-300 bp is in the 1.1X sample, the 0.8X sample was kept in case more DNA was needed. DNA from both samples 0.8X and 1.1X were quantified by running 1 µL on a 2% agarose gel along with different amounts of low molecular weight DNA ladder (100 ng, 200 ng, 400 ng).

**End Repair:** 50 µL of Hi-C sample was transferred to a PCR tube, then 20 µL of the end-repair mix [3.5X NEB ligation buffer (NEB B0202S), 17.5 mM dNTP mix, 7.5 U T4 DNA polymerase (NEBM0203L), 25 U T4 polynucleotide kinase (NEB M0201S), 2.5 U Klenow polymerase Polymerase I (NEB M0210L)] was added. The 70 µL total volume reaction was then incubated at 37°C for 30 min, followed by incubation at 65°C for 20 min to inactivate Klenow polymerase, and then the sample was put at 4°C. The volume was brought up to 400 µL using 1X TLE for the next step.

**Biotin pull-down:** All the following steps were performed with 1.5 mL loBind tubes (Eppendorf 22431021). 15 µL of MyOne streptavidin C1 beads mix (Thermo Fisher 65001) was transferred to a 1.5 mL tube. The beads were washed twice by adding 400 µL of TWB [5 mM Tris-HCl pH8.0, 0.5 mM EDTA, 1 M NaCl, 0.05% Tween20] followed by incubation for 3 min at RT. The tube was then placed on an MPS for 1 min and the supernatant was removed. After the washes, the beads were resuspended in 400 µL of 2X Binding Buffer (BB) [10 mM Tris-HCl pH8, 1 mM EDTA, 2 M NaCl] and mixed with the 400 µL DNA from the previous step in a new 1.5 mL tube. The mixture was incubated for 15 min at RT with rotation, the tube was then placed on the MPS for 1 min and the supernatant was removed. The DNA bound to the beads was washed by adding 400 µL of 1X BB and transferred to a new tube. The beads were reclaimed against the MPS for 1 min, and the supernatant was discarded. The second wash used 100 µL of 1X TLE, beads were reclaimed against MPS for 1 min, and the supernatant was discarded. Finally, the DNA bound to the beads was eluted in 32 µL of 1X TLE.

**A-tailing:** A dATP was added to the 3’ ends by adding 18 µL of A-tailing mix [5 µL NEB buffer 3.1, 10 µL of 1 mM dATP, 3 U Klenow exo (NEB M0212S)] to the 32 µL of DNA sample from the previous step. The reaction was incubated in a PCR machine [at 37°C for 30 min, then at 65°C for 20 min, followed by cool down to 4°C]. Next, the tube was placed on ice immediately. The sample was transferred to a 1.5 mL loBind tube, the tube was placed on the MPS for 1 min and the supernatant was removed. The streptavidin beads bound to DNA were washed twice using 100 µL 1X T4 DNA Ligase Buffer (Invitrogen). Finally, streptavidin beads bound to DNA were resuspended in 40 µL 1X T4 DNA Ligase buffer (Invitrogen).

**Illumina adapter ligation and paired-end PCR**: For this step, the TruSeq DNA LT kit Set A (REF#15041757) was used. 10 µL of ligation mix [5 µL Illumina paired-end adapters, 3 µL T4 DNA ligase Invitrogen, 2 µL 5x T4 DNA ligase buffer (Invitrogen 5X)] was added to the 40 µL Hi-C sample from the previous step. The ligation sample was then incubated at RT for 2 hours on a rotator. The sample was transferred to a 1.5 mL loBind tube, the tube was placed on the MPS for 1 min and the supernatant was removed. The streptavidin beads bound to DNA were washed twice with 400 µl of **TWB, then twice** using 100 µL 1X TLE. Finally, the sample was resuspended in 20 µL 0f 1XTLE.

**Illumina Truseq Kit for PCR:** We performed three trial PCR reactions as follows [2.5 µL DNA bound to beads, 2 µL of Primers mix (TruSeq DNA LT kit Set A 15041757)), 10 µL Master Mix (TruSeq DNA LT kit Set A 15041757), 10.5 µL of ultrapure distilled water (Invitrogen)]. We split the 25 µL over three PCR tubes (5 µL, 5 µL, 15 µL per tube). Each of the three samples was then amplified with different numbers of PCR cycles (6, 8, 10 respectively) to assess the Hi-C library quality: [30 sec at 98°C, n cycles of (30 sec at 98°C, 30 sec at 65°C, 30 sec at 72°C), 5 min at 72°C, hold at 10°C]. 10 µL was taken from the 15 µL sample (with 10 PCR cycles), the 10 µL sample was then digested with ClaI for 1 h by adding 10 µL of digestion mix [1.5 µL **10x NEB Cutsmart buffer, 1.5 µL ClaI** (NEB R0197S), 7 µL ultrapure distilled water]. The 5 µL of each PCR cycle sample along with the 20 µL digested sample, and titration of the low molecular ladder (100 ng, 200 ng, 400 ng) (NEB) were run on a 2% Agarose gel. After digestion with ClaI, a downward shift of the amplified DNA to smaller sizes is expected, which indicates DNA ends were correctly filled in and ligated (creating a ClaI site). The number of PCR cycles to generate the final Hi-C material for deep sequencing was chosen based the minimum number of PCR cycles in the PCR titration that was needed to obtain sufficient amounts of DNA for sequencing using the remaining 17.5 µL Hi-C sample.

### DpnII-Seq

For each DpnII-Seq library, 10 million nuclei were used right after the pre-digestion procedure described above (Pre-digestion of nuclei). The pre-digested nuclei were then treated as follows:

**Proteinase K**: 50 µL of 10 mg/mL proteinase K (ThermoFisher # 25530) was added to each 500 µL pre-digested nuclei sample (2 million nuclei) (See Methods: Pre-digestion) and the 5 tubes were incubated at 65°C for 3 hours.

**DNA purification**: Tubes were cooled to room temperature and all 5 samples were pooled in a single 15 mL tube (2.75 mL total volume). The DNA was extracted by adding an equal volume of 2.75 mL of saturated phenol pH8.0: chloroform (1:1) (Fisher BP1750I-400), followed by vortexing for 1 min. The sample (5.5 mL) was transferred to a 15 mL phase-lock tube (Quiagen #129065) followed by centriguation at 5,000 g for 10 min. The upper phase was transferred to a 15 mL tube to start the second extraction. An equal volume of 2.75 mL saturated phenol pH8.0: chloroform (1:1) was added, followed by vortexing for 1 min. Then the mix was transferred to a 15 mL phase-lock tube (Quiagen #129065) followed by centrifugation at 5,000 g for 10 min. The upper phase of ∼ 2.75 mL was transferred to a 15 mL tube (high speed), 1/10 volume (275 µL) 3M sodium acetate pH 5.2 was added followed by brief vortexing and then 2.5 volumes of ice-cold 100% ethanol (6.875 mL) were added. The tube was inverted slowly several times, incubated at −80°C for 1 hr and then DNA was pelleted by centrifugation at 16,000 g for 30 min at 4°C. The supernatant was discarded and the pellet was dissolved in 500 µL 1X NEB3.1 and transferred to a 0.5 mL AMICON Ultra Centrifuge filter (UFC5030BK EMD Millipore). The column was centrifuged for 5 min at 14,000 g and the flow-through was discarded. The column was washed 4 times using 450 µL of 1X NEB3.1 for desalting of DNA. After the final wash, the library remaining in the column (∼50 µL) was eluted in 450 µL of 1XNEB3.1; the column was flipped upside down into a new tube to collect DNA and centrifuged for 3 min at 4,000 g. ∼500 µL of DNA was recovered. RNA was degraded by adding 1 µL of 10 mg/mL RNAase A and incubation for 30 min at 37°C. The amount of DNA was estimated by running an aliquot on a 1% Agarose gel along with a 1kb ladder (NEB#N3232s).

**Biotin Fill-in**: 1XNEB3.1 was added the reaction to a final volume of 680 µL, and then the 680 µL was split over 2 1.5 mL tubes. DNA ends were filled in and marked with biotin-14-dATP. To each tube 60 µL of biotin fill-in master mix was added: [1xNEB2.1, 0.25 mM dCTP, 0.25 mM dGTP, 0.25 mM dTTP, 0.25 mM biotin-dATP (ThermoFisher#19524016), 50 U Klenow polymerase Polymerase I (NEB M0210L)]. Samples were incubated at 37°C in a Thermocycler for 75 mins. Next, the tubes were placed on ice immediately for 15 mins, and samples from the 2 tubes were combined to obtain a final volume ∼800 µL. Amicon filters were used to reduce the volume of the final sample from 801 µL to 130 µL.

**Sonication**: DNA was sonicated to a size of 100 – 300 bp using a Covaris instrument (Duty Cycle 10%, Intensity 5, Cycles per Burst 200, set Mode Frequency sweeping, continuous degassing, process time 60 sec, Number of cycles) for 4 cycles. The 130 µL of sonicated DNA was transferred to a 1.5 mL tube and 1XTLE was added to a total volume of 500 µL. DNA fragment size was determined by running 2 µL of DNA along with low molecular ladder (NEB) on a 2% agarose gel.

**Size fractionation using AMpure XP:** 500 µL AMpure beads (Beckman Coulter A63881) were added to a 1.5 mL tube labeled as 1.1X. The tube was placed on the MPS for 5 min, and the supernatant was removed. Beads were resuspended in 150 µL AMpure mixture in order to make 1.1X. 400 µL of AMpure mixture was added to 500 µL of sonicated DNA from the previous step and the tube was labeled 0.8X. The sample was vortexed and centrifuged briefly using a tabletop small centrifuge followed by incubation at RT for 10 min on a rocking platform. Then the tube was placed on the MPS for 5 min at RT. The 0.8X supernatants were collected and added to the 1.1X tube, the tube was briefly vortexed and centrifuged followed by incubation at RT for 10 min on a rocking platform. The tube was placed on the MPS for 5 min at RT, and the supernatant discarded. Beads in the 0.8X and 1.1X tubes were washed twice with 1 mL 70% ethanol, reclaiming beads against the MPS for 5 min. Beads on the MPS were then dried until ethanol had evaporated completely. Next, 51 µL of 1XTLE was added to the 0.8X and 1.1X tubes to elute DNA from the beads. Tubes were incubated at RT on a rocking platform for 10 min. The 0.8X and 1.1X tubes were placed on the MPS for 5 min. Finally, the supernatants were transferred to 1.7 mL tubes labeled 0.8X and 1.1X. The 1.1X sample contains DNA that ranges in size from 100-300 bp. The DNA in the 0.8X sample was kept in case more DNA was required, in which case the DNA would be sonicated using 2 cycles followed by a similar round of size fractionation as described above. The amount of DNA from both samples 0.8X and 1.1X was quantified by running 1 µL on a 2% agarose gel along with a titration of low molecular weight DNA ladder (100 ng, 200 ng, 400 ng).

**End Repair:** 50 µL from the 1.1X sample was transferred to a PCR tube, and 20 µL of end repair mix was added: [3.5X NEB ligation buffer (NEB B0202S), 0.875 mM dNTP mix, 0.375 U/µL T4 DNA polymerase, 1.25 U/µL T4 polynucleotide kinase, 0.125 U/µL Klenow DNA polymerase]. The 70 µl total volume reaction was incubated for 30 min at 20°C in a PCR machine and then placed on ice. The DNA was purified by 1:2 Ampure, by adding 140 µL 2X Ampure solution to the 70 µL DNA sample followed by incubation for 5 min at RT. The tube was placed on the MPS for 4 min to reclaim the beads and the supernatant was discarded. The beads were washed twice with 1 mL of 70% ethanol while on the MPS. After beads were dried DNA was eluted in 32 µL TLE (pH 8.0) and incubation for 10 min at RT. The supernatant was transferred to a 1.5 mL tube.

**A-tailing:** A dATP was added to the 3’ ends by adding 18 µL of A-tailing mix [5 µL NEB buffer 3.1, 10 µL of 1 mM dATP, 3 U Klenow exo (NEB M0212S)] to the 32 µL of DNA sample from the previous step. The reaction was then incubated in a PCR machine at 37°C for 30 min followed by incubation 65°C for 20 min and cooling down to 4°C. The tube was placed on ice. The volume was brought to 100 µL by adding 1X NEB2.1. The DNA was then purified by adding 1:2 Ampure mix (200 µL of Ampure was added to the 100 µL final DNA volume). Finally, the DNA was eluted in 40 µL of 1X T4 DNA ligase buffer (Invitrogen 5X).

**Illumina adapter ligation and paired-end PCR**: For this step we used the TruSeq DNA LT kit Set A (REF#15041757). 50 µL of ligation mix [25 µL Illumina paired-end adapters, 15 µL T4 DNA ligase Invitrogen, 10 µL 5X T4 DNA ligase buffer (Invitrogen 5X)] was added to the 40 µL sample from the previous step. The ligation sample was then incubated at RT for 2 hours on a rotator. Next, the DNA was purified by adding 1:1 Ampure solution (180 µL of Ampure mix was added to the 90 µL sample), the supernatant was discarded and beads were washed twice with 1 mL of 70% ethanol. After the last wash step, the beads were resuspended in 400 µL of 1X TLE and incubated at RT on a rocking platform for 10 mins. The tube was placed on the MPS for 4 mins. Finally, the 400 µL supernatant was transferred to a new tube.

**Biotin pull-down**: All the following steps are done using 1.5 mL loBind tube (Eppendorf 22431021). 15 µL of MyOne streptavidin C1 beads mix (Thermo Fisher 65001) was transferred to a 1.5 mL tube. The beads were washed twice with 400 µL of TWB [5 mM Tris-HCl pH8.0, 0.5 mM EDTA, 1 M NaCl, 0.05% Tween20] by incubation for 3 min at RT. After each wash, the tube was placed on the MPS for 1 min and the supernatant was removed. After the washes, the beads were resuspended in 400 µL of 2X Binding Buffer (BB) [10 mM Tris-HCl pH8.0, 1 mM EDTA, 2 M NaCl] and mixed with the 400 µL DNA from the previous step in a new 1.5 mL. The mixture was incubated for 15 min at RT with rotation. The tube was then placed on the MPS for 1 min and the supernatant was removed. The DNA bound to the beads was washed first by adding 400 µL of 1X BB and transferring to a new tube. The beads were reclaimed against the MPS for 1 min, and the supernatant discarded. 100 µL of 1X TLE was added and the beads were reclaimed against the MPS for 1 min, then the supernatant was discarded. Finally, the DNA bound to the beads was eluted in 32.5 µL of 1X TLE.

**PCR optimization:** The Illumina Truseq Kit (DNA LT kit Set A (REF#15041757)) was used for PCR amplification of DNA for DpnII-Seq. The trial PCR reaction was set up as follows: [2.5 µL DNA bound to beads, 2 µL of Primers mix (Truseq kit), 10 µL Master Mix (Truseq kit), 10.5 µL of ultrapure distilled water (Invitrogen)]. The 25 µL was split over four PCR tubes (5 µL/per tube). Each of the four samples was incubated for different PCR cycles (6, 8, 10, or 12 cycles): [30 sec at 98°C, n cycles of (30 sec at 98°C, 30 sec at 65°C, 30 sec at 72°C), 7 min at 72°C, hold at 10°C]. The optimal PCR cycle number needed to get enough DNA for sequencing was determined by running the 4 PCR reactions on a 2% agarose gel along with low molecular ladder titration (100 ng, 200 ng, 400 ng). Three PCR reactions of 50 µL volume were then performed: [5 µL DNA bound to beads, 4 µL of Primers mix (Truseq kit), 20 µL Master Mix (Truseq kit), 21 µL of ultrapure distilled water (Invitrogen)]. The 3 PCR reactions were pooled together to obtain 150 µL total volume. The samples were reclaimed against the MPS for 1 min, then the PCR products (supernatant) were taken to new 1.5 mL tubes. 1:1 Ampure was performed for removal of primer dimers (150 µL of Ampure and 150 µL DNA sample). Finally, beads were resuspended in 35 µL of TLE to elute the DNA. DNA that remained bound to beads was saved after a first wash using TBW followed by two washes with 1X TLE and then resuspended in 30 µL of 1X TLE.

### Lamin A Immunofluorescence and DAPI

For nuclei immunofluorescence, we prepared a coverslip by adding 1 mL of 0.1% Poly-L-lysine solution (Sigma SLBQ5716V) for 10 min, then coverslips were dried using Whatman papers. Each coverslip was transferred to a single well of an eight wells plate. The coverslips were washed twice using PBS. Next 500 µL of 30% sucrose with 1 mM DTT was added on top of the coverslips to protect nuclei from an abrupt contact with coverslip during spinning. 1 million control nuclei or nuclei after chromatin digestion were crosslinked for 20 min using a 4% final concentration of Paraformaldehyde immediately after pre-digestion. Next, nuclei were added slowly on top of he sucrose solutions on the coverslips and spun for 15 mins at 2,500 g at 4°C. Next, nuclei were assumed to be attached to the coverslips which were then transferred to a new 8 well plate. The coverslips were washed five times with 1% PBS. Next, non-specific binding of the primary antibody was blocked by adding 500 µL of the blocking buffer [3% BSA, 1X PBS, 0.1% Triton X-100 (Sigma <u>9002-93-1</u>)] and incubating for 60 min at RT. Afterward, lamin A antibody (ab 26300) was diluted 1:1000 in blocking buffer, and the coverslip was incubated face-down on top of a 250 µL of lamin A antibody droplet that was placed on parafilm for 120 min at RT. Then, the coverslip was placed back in the well of a new plate face-up and washed five times with washing buffer (1X PBS, 0.1% Triton X-100). The secondary antibody Goat Anti-Rabbit (ab150077) was diluted 1:1000 in blocking buffer, and the coverslip was incubated face-down on top of a 250 µL droplet of the secondary antibody (Goat Abti-Rabbit (ab150077) that was placed on parafilm for 60 min at RT. Next, the coverslip was placed back in the well of a new plate face-up and washed five times with washing buffer (1X PBS, 0.1% Triton X-100) and twice with 1X PBS. The slide was mounted and sealed using 10 µL antifade mountant with DAPI (Invitrogen P36931).

For image acquisition, we used a Nikon Eclipse Ti microscope. Imaging was performed using an Apo TIRF, N.A. 1.49, 60X oil immersion objective (Nikon), and a Zyla sCMOS camera (Andor). Z-series of 0.2 µm slices were acquired using Nikon Elements software (Version 4.4).

### Chromatin fractionation assay

Chromatin-bound proteins were isolated and separated from free proteins. A sample of 2 million control nuclei or pre-digested nuclei (obtained as described above “Pre-digestion of nuclei”) was centrifuged at 5,000 g for 5 min at 4°C. The supernatant was transferred to an Amicon column to reduce the volume from 500 µL to 100 µL by centrifugation for 4 min at 14,000 g. This sample contains the free protein fraction. Next, 26 µL of glycerol and 1.3 µL of 100X protease inhibitor cocktail were added to the 100 µL free proteins sample. The pellet containing the nuclei was resuspended in 100 µL of nuclei purification buffer with Triton (10 mM PIPES pH 7.4, 10 mM KCl, 2 mM MgCl2, 0.25% Triton, 1% Protease inhibitor, 1mM DTT) and incubated for 10 min on ice. Then, in order to protect protein structure during sonication, 25 µL of glycerol was added to the 100 µL pellet sample to have 20% final glycerol concentration. The sample was sonicated using a Covaris instrument at 4°C as follows: (Duty Cycle 10%, Intensity 5, Cycles per Burst 200, set Mode Frequency sweeping, continuous degassing, process time 60 sec, 4 cycles). The pellet sample contains chromatin-bound proteins, was transferred to a 1.5 mL tube. All samples were stored at −20. These samples contain the protein bound CTCF and cohesin. Note: when these samples were centrifuged after the triton solubilization, we found that no SMC3 or CTCF could be detected in the supernatant. These results indicate that non-chromatin-bound proteins exit the nuclei and were recovered in the supernatant prior to triton solubilization step.

For analysis of CTCF and SMC1 chromatin binding: 15 µL from each protein sample (supernatant or pellet) was mixed with 5 µL of 5X Lane Marker Reducing Sample Buffer (Thermo Fisher 39000), then the mix was boiled for 10 min. The samples were cooled down to RT before loading them on a 3-8% Tris-Acetate Protein Gels (Invitrogen EA0375PK2). Next, the gel was run in 1X Tris-Acetate SDS Running Buffer (Invitrogen LA0041) for 75 min at 150V. For Histone H3: 1 µL of protein sample was mixed with 14 µL of PBS containing 1% Protease inhibitor, 5 µL of 5X Lane Marker Reducing Sample Buffer was added to the mix and boiled for 10 min. The samples were cooled down to RT before loading them in Tris-Base 4-12% (Invitrogen NP0322BOX), then the gel was run in 1X MES-SDS running buffer (Invitrogen B0002) for 60 min at 150V. The proteins were transferred from the gel to nitrocellulose membrane using 1X western blot transfer buffer (Thermo science 35040). The transfer was 120 min for SMC1 and CTCF and 75 min for H3. The nitrocellulose membranes were washed using 1X TBST [50 mM Tris-Cl, pH 7.6; 150 mM NaCl, 0.1 mL of Tween 20], then Blocked for 120 min using 5% milk (1 g milk in 20 mL 1X TBST). The membrane when then incubated overnight at 4°C with primary antibody diluted in 5% milk [1:1000 CTCF antibody cell signaling (activeMotif 61311), 1:2000 SMC1 (Bethyl Antibody, A300-055A), 1:4000 H3 Abcam (ab1791)]. Next, the membranes were washed 6 times for 10 min per wash using 1X TBST. The secondary antibody anti-rabbit IgG HRP from cell signaling was diluted using 5% milk for CTCF and SMC1 [1:1000 for CTCF, 1:2000 SMC1] and in 1% milk for H3 1:5000 dilution. Membranes were incubated for 120 min at RT. Finally, membranes were washed 6 times for 10 min using 1X TBST. Finally, the membranes were developed using luminol-based enhanced chemiluminescence(Thermo science 34076).

### Micromanipulation force measurement and treatments of an isolated nuclei

Micromanipulation force measurements were conducted as described previously in Stephens et al. (Stephens et al., 2017). K562 cells were grown in microscope slide wells and treated with 1 µg/mL latrunculin A (Enzo Life Sciences) for ∼45 min before single nucleus isolation. The nucleus was isolated by using small amounts of detergent (0.05% Triton X-100 in PBS) locally sprayed onto a living cell via an “isolation” micropipette. This gentle lysis allows the use of a second micropipette to retrieve the nucleus from the cell, using slight aspiration and non-specific adherence to the inside of the micropipette. A third micropipette was then attached to the opposite end of the nucleus in a similar fashion. This last “force” micropipette was pre-calibrated for its deflection spring constant, which is on the order of 2 nN/µm. A custom computer program written in LabView was then run to move the “pull” micropipette and track the position of both the “pull” and “force” pipettes. The “pull” pipette was instructed to move 5 µm at 45 nm/sec. The program then tracked the distance between the pipettes to provide a measure of nucleus extension ∼3 µm. Tracking the distance that the “force” pipette moved/deflected multiplied by the pre-measured spring constant provides a calculation of force exerted. Calculations were done in Excel (Microsoft) to produce a force-extension plot from which the best-fit slope of the line would provide a spring constant of the nucleus (nN/µm). Isolated nuclei were measured twice initially to establish the native spring constant prior to treatment. After 50 uL of buffer only (control), 100 units DpnII (|GATC) with NEB buffer 3.1, or 100 units HindIII (A|AGCTT) with NEB buffer 2.1 was added to the 1.5 mL imaging well and mixed gently. Force measurements were performed 5 min, 30 min, and 60 min post-treatment.

## QUANTIFICATION AND STATISTICAL ANALYSIS

### 3C-PCR

The human β-globin locus is an ideal region to examine looping interactions between enhancers and genes because of the strong looping interactions between the LCR and HBG globin gene in the erythroleukemia cell line K562, which highly expresses the globin genes (Dostie et al., 2006). 3C libraries were generated from: (1) K562 cells that have an LCR-HBG interaction, (2) GM12878 cells in which the LCR-HBG looping interaction is absent, and (3) beta-globin BAC (ENm009) control to normalize for primer bias. To investigate the interaction between the LCR and HBG gene, 3C primers from (Dostie et al., 2006) were used. 16 forward primers of 28-33 bp length were designed 40-60 bp upstream of each EcoRI site throughout a 110 kb region around the Beta Globin locus (chr11: 5221788-5337325). The EcoRI fragment overlapping with the LCR (HS3,4,5) was used as an anchor to detect the interaction frequencies between the LCR and EcoRI fragments throughout the β-globin locus. For each primer pair, triplicate PCR reactions were set up, and the mean of the three was normalized to the BAC signal for the same primer pair before plotting normalized interaction frequency in the y-axis, the distance from EcoRI fragment overlapping with LCR to neighboring EcoRI fragments is plotted in the x-axis. Error bars are the standard error of the mean (SEM). PCR primers are listed in Supplemental Table S1.

### 5C data processing

The fastq files for 5C sequencing data were processed as described in https://github.com/dekkerlab/5C-CBFb-SMMHC-Inhib/blob/master/data_processing_steps.md The Fastq files were mapped using novoalign to a reference genome built from the pool of all 277 probes. After mapping, we combined the read-pairs. The results were then transferred to a matrix format, and interactions were filtered as previously described (Lajoie et al., 2009; Sanyal et al., 2012). First, interactions that belong to the same EcoRI fragment were removed. Second, outliers that are overrepresented as a result of overamplification were also removed. Outliers were defined as the interactions with a Z-score greater than 20 in all datasets. Third, probes that strongly over or underperform which leads to strongly enriched or depleted interactions in a whole row of interactions, were also removed. The four matrices were then scaled to the same number of total reads. Finally, data were binned at 20 Kb (median) with a sliding window with 2.5 Kb steps.

### Hi-C data processing

Hi-C read mapping, filtering, binning and matrix normalization were performed using the cMapping pipeline available at https://github.com/dekkerlab/cMapping (Lajoie et al., 2015). In brief, Hi-C reads were mapped to reference human genome assembly hg19 using an iterative mapping strategy and Bowtie 2 (Langmead et al., 2009). Successfully mapped reads were then filtered to remove reads mapping to the same restriction fragment and to remove PCR duplicates. Interaction frequency versus distance plots displayed high variance for interactions - below 1 kb for all samples. Hence, after mapping of valid pair, we removed all pairs with a genomic distance less than 1 kb. The remaining valid read pairs were then binned to 500 kb, 40 kb, and 10 kb resolution matrices. Outlier bins of these matrices with low signal were assigned values of NA. Then as a bias correction step, matrices were normalized such that the sum of interactions in each row/column are approximately equivalent via an iterative correction procedure (ICE) (Imakaev et al., 2012). Lastlly, for comparison between samples, matrices were scaled such that the total interactions for a genome-wide matrix equals one billion for each sample. These ICEd scaled matrices were used for subsequent analyses.

### A/B compartments

All reads from Hi-C in control K562 samples were pooled to identify A (active) and B (inactive) compartments in K562 cells. A/B compartments were identified at 40 kb resolution following the procedure described in (Lieberman-Aiden et al., 2009) using matrix2compartment.pl in https://github.com/dekkerlab/cworld-dekker. Briefly, each *cis* interaction matrix was first transformed into a z-score matrix followed by transformation into a correlation matrix. PCA was performed on the correlation matrix and the first eigenvector (PC1) of the PCA analysis was used to identify compartments for each chromosome. A/B compartments were assigned based on gene density such that the A-compartment was more gene-dense than the B-compartment. Positive PC1 values indicate gene-rich A compartments and negative PC1 values indicate gene-poor B compartments. For chromosome 9 the compartments were called for each chromosome arm separately as PC1 captured preferences for interactions within the same arm as opposed to canonical compartment preferences.

### LOS and half-life calculation

To measure the 3D structure changes resulting from DpnII or HindIII pre-digestion we quantified the amount of cis interactions lost or gained in a 6 Mb window centered at every 40 kb bin genome wide. For each 40 kb bin, the percent of interactions occurring within its 6 Mb window (corresponding to interactions less than or equal to 3 Mb in distance either upstream or downstream from 40 kb bin) out of total interactions for the 40 kb bin (cis and trans) was calculated. These 6 Mb cis percentages were calculated for control, DpnII pre-digested, and HindIII-pre-digested nuclei. The change in 3D structure relative to control using these cis percentages was given by the following loss of structure (LOS) metric:

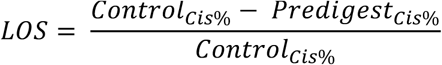

Hence, LOS values in the range (0, 1) represent a loss in short range contacts after pre-digestion; LOS values < 0 represent an increase in short range contacts after pre-digest, and an LOS equal to zero would indicate no change in structure after pre-digestion. A window of 6 Mb was chosen as we sought here to quantify interactions disrupted by pre-digestion. Many longer range interactions increased after pre-digestion, potentially due to random ligations of cut fragments that start to mix. Differences noted in A and B stability was preserved when LOS was calculated using cis percents for entire chromosomes as opposed to a 6 Mb window, however the size of chromosomes did bias results by giving 40 kb bins in small chromosomes greater LOS.

To quantify the timing of disrupted interactions we generated a half-life track utilizing the Hi-C matrices from the DpnII timecourses. For each 40 kb bin we fit a curve to the LOS of each timepoint following an exponential decay of the form (Supplemental Figure 4C):

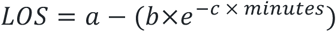

such that *a*, *b* and *c* are parameters to fit.The half-life, t_1/2_, was defined as the time required to reach half saturation, saturation being the 16 hour timepoint where maximal cis interactions have been lost. Half-life values were then computed for every 40 kb bin genome wide. To remove noisy and less reliable t_1/2_ data, we first removed all extreme outliers bins where the sum of squared residuals (SSR) for the exponential fit was greater than 0.1. Then all bins with an SSR greater than two standard deviations from the mean was deemed an outlier and also removed from analyses.

As LOS and t_1/2_ are both dependent on digestion efficiency we also generated residual LOS and t_1/2_ tracks to account for bin to bin variation in digestion efficiency. We used a moving average approach to calculate residuals for LOS as a function of DpnII-seq signal and also t_1/2_ as a function of DpnII-seq signal since the relationships between these variables were non-linear (Figure 3F left, Supplementary Figure 5C). For both stability metrics LOS and t_1/2_, a sliding window of 200 DpnII-seq signal with a step size of one was used to calculate mean LOS or t_1/2_ signal for each DpnII-seq signal increment (Figure 3F left, Supplementary Figure 5C). Window and step size were selected by manual inspection of moving averages and compromising between over and underfitting. These moving averages were used to calculate residuals such that a positive LOS residual indicates more structure loss than expected by given digestion efficiency and a negative LOS residual indicates less structure loss than expected. As t_1/2_ is inversely related to LOS, positive t_1/2_ residuals indicate less structure loss than expected and negative t_1/2_ residuals indicate more structure loss than expected. Moving averages were also used to generate residuals for DpnII-seq as a function of PC1 and LOS as a function of PC1 (Figure 3G, right)

### DpnII-seq data analysis

Sequenced reads were mapped to the genome using the Bowtie read aligner (Langmead et al., 2009) and reads mapping to multiple sites of the genome were removed. As expected, a high percentage of reads mapped precisely to their associated restriction cut site (Supplementary Figure 3C). To remove potential artificial biases, we filtered out paired-end reads from fragments whose start or end coordinate was more than three nucleotides from an appropriate restriction cut site. Filtered reads were then binned to 500 kb or 40 kb resolutions. The K562 cell line has a primarily triploid karyotype with regions of the genome in diploid and tetraploid states. Copy number state assignments for each 500 kb or 40 kb bin were assigned using publicly available K562 copy number data from the Catalogue of Somatic Mutations In Cancer (COSMIC) database (https://cancer.sanger.ac.uk/cell_lines/download). Copy number segments in the COSMIC dataset were identified by PICNIC analysis of Affymetrix SNP6.0 array data (PMID:19837654). Read coverage files at 500 kb and 40 kb were corrected to a genome wide diploid state using the copy number state assignments and dividing coverage by appropriate correction value (diploid = 1, triploid =1.5, tetraploid = 2, etc.) per bin. (Supplementary Figure 3D,E). Final copy number corrected coverage files were used for all downstream analysis and are available (Supplementary File). DpnII-seq computational workflow is maintained at https://github.com/tborrman/DpnII-seq

### Subcompartments

Rao et al. (2014) divided the canonical A/B compartments into five primary subcompartments A1, A2, B1, B2, B3 based on each subcompartment’s preferential Hi-C interactions in GM12878 cells. Subcompartments were annotated using high resolution (∼1 kb) Hi-C data and were shown to display unique genomic and epigenomic profiles. K562 subcompartments were annotated in (Xiong and Ma, 2019) via the method SNIPER using lower resolution Hi-C data. In short, SNIPER infers subcompartments via a neural network approach to accurately annotate subcompartments using Hi-C datasets with moderate coverage (∼500 million mapped read pairs). Xiong et al.’s K562 SNIPER subcompartments showed a substantial conservation with GM12878 annotations from Rao et al. (Rao et al., 2014) and were also enriched in similar epigenetic features, hence we utilized these SNIPER annotations to compare subcompartment status with chromatin stability. K562 SNIPER subcompartments were annotated at 100 kb resolution. To compare with our 40 kb resolution liquid chromatin Hi-C data, we binned the 100 kb subcompartment annotations to 40kb such that any 40 kb bin overlapping a boundary of two separate subcompartments was assigned a value of NA. Upon piling up K562 subcompartment boundaries, we also found enrichment and depletion of various chromatin features consistent with those described in both Rao et al. (Rao et al., 2014) and Xiong et al. (Xiong and Ma, 2019).

### Sub-nuclear structures

To assess the effect of sub-nuclear structures on chromatin stability we utilized the extensive genetic and epigenetic data publicly available for K562 cells (Supplemental Table S3).

Fold change over control ChIP-seq tracks for histone modifications, chromatin remodellers, and other various proteins were downloaded from the ENCODE Portal. To compare ChIP-seq data with t_1/2_, or residuals of t_1/2_ after correction for DpnII-signal, we binned the ChIP-seq signal tracks into 40 kb such that each 40 kb bin represented the mean signal found across the bin. Bins with no overlapping signal were designated a value of NA.

To examine the association between methylation state and t_1/2_ or residuals of t_1/2_ after correction for DpnII-signal, we downloaded methylation state at CpG Whole-Genome Bisulfite Sequencing (WGBS) tracks from ENCODE. As the methylation data was mapped to hg38, we used the UCSC LiftOver program to convert coordinates to hg19. Then percentage methylation at CpG sites was binned to 40 kb resolution using the mean.

As there is currently no nucleolus associated domains (NADs) data available for K562, we analyzed a binary NADs state track for the human embryonic fibroblast IMR90 cell line (Dillinger et al., 2017). Dillinger et al. annotated NADs via a two-state hidden Markov model of aCGH data from DNA of isolated nucleoli. Using these annotated NADs, coverage of each 40 kb bin for NADs was assessed and used for all our downstream analyses.

Mapping of nuclear speckle, nuclear lamina and PolII associated loci for K562 cells was accomplished recently via the TSA-seq protocol (Chen et al., 2018). Signal tracks of log2(pull-down/input) were downloaded from GEO and binned to 40 kb as previously described for ChIP-seq files. Microarray data for LaminB1 associated domains identified through the DamID protocol was also available from that study. We used the UCSC LiftOver program to convert coordinates from hg18 to hg19. We then binned the log2(Dam-LaminB1/Dam) signal to 40 kb bins as previously described for ChIP-seq files.

To analyze cell cycle relationship with chromatin stability we downloaded Repli-seq data for K562 cells from ENCODE. Actively replicating regions are quantified as a percentage normalized signal for FACS sorted cells in G1 phase, four stages of S phase (S1-S4) and G2 phase. Signal tracks for Repli-seq data were binned to 40 kb as previously described for ChIP-seq files.

Binning of data was performed using the bedtools/v2.26.0 software. To assess the quality of the publicly downloaded data we generated the spearman correlation matrix of all binned signal tracks (Figure S6A). Hierarchical clustering of rows of the correlation matrix position heterochromatic marks (H3K9me3, HP1α, HP1β, NADs, and LADs) near one another as expected. The majority active marks form a larger cluster, with the markers for polycomb regions (H3K27me3, CBX8, BMI1, RNF2, and SUZ12) representing facultative heterochromatin clustered together segregating active from inactive marks.

### Gene Expression

To assess the effect of gene expression on chromatin stability we utilized processed gene expression quantifications of total RNA-seq for K562 cells available from ENCODE (Accession ID: ENCFF782PCD). Gene locations were mapped using the hg19 ensGene table from UCSC Table Browser. To compare expression values with 40 kb resolution t_1/2_ or residuals of t_1/2_ after correction for DpnII-signal tracks, fragments per kilobase million (FPKM) values for each gene were binned to 40 kb such that each 40 kb bin represented the mean FPKM for all genes overlapping that bin. Bins without any genes were assigned a value of NA. Binned FPKM >=1 was determined to be a reasonable cutoff for expression by inspection of the full distribution of FPKM values.

### Compartmentalization saddle plots

Saddle plot was adapted from cool tools (https://github.com/hms-dbmi/hic-data-analysis-bootcamp/tree/master/notebooks/04_analysis_cooltools-eigenvector-saddle.ipynb). To measure the strength of compartments the average intra-chromosomal interactions frequencies between 40 kb bins were normalized by genomic distance (observed/expected Hi-C maps). Then, in a 50 by 50 bin matrix, the distance corrected interaction bins were sorted based on their PC1 value in increasing order. Finally, all intra-chromosomal interactions with similar PC1 values were aggregated to obtain compartmentalization saddle plots. In these plots, preferential B-B interactions are in the upper left corner, and preferential A-A interactions are in the lower right corner.

#### Homotypic interaction saddle plots

The average intra-chromosomal interactions frequencies between 40 kb bins were normalized by genomic distance (observed/expected Hi-C maps). Then, in a 50 by 50 bin matrix, the distance corrected interaction bins were sorted based on their signal value (TSA-seq, DamID) for a given factor (SON, Lamin). Finally, all intra-chromosomal interactions with similar signal values were aggregated to obtain homotypic interaction saddle plots. In these plots, pair-wise interactions between loci enriched in factor binding are shown in the lower right corner, and pair-wise interactions between loci not bound by the factor are shown in the upper left corner.

### Scaling plot

The script to generate scaling plots was adapted from cooltools (https://github.com/mirnylab/cooltools/tree/master/cooltools). Genome-wide curves of normalized contact frequency *P*(*s*) is plotted as a function of genomic distance for all intra-chromosomal interactions. Each library was normalized by total valid interactions

### Mean z-score heatmap

Each genome wide 40kb signal vector for a sub-nuclear structure was cleaned for outliers above three standard deviations of the vector’s mean. Each cleaned vector was z-score transformed and then partitioned based on the different t_1/2_ residual intervals for associated bins. The mean z-score for all bins within a given t_1/2_ residual interval is plotted as a square in the heatmap.

