## Supplemental Figures for "Compartment-dependent chromatin interaction dynamics revealed by liquid chromatin Hi-C"

Supplemental Figure 1

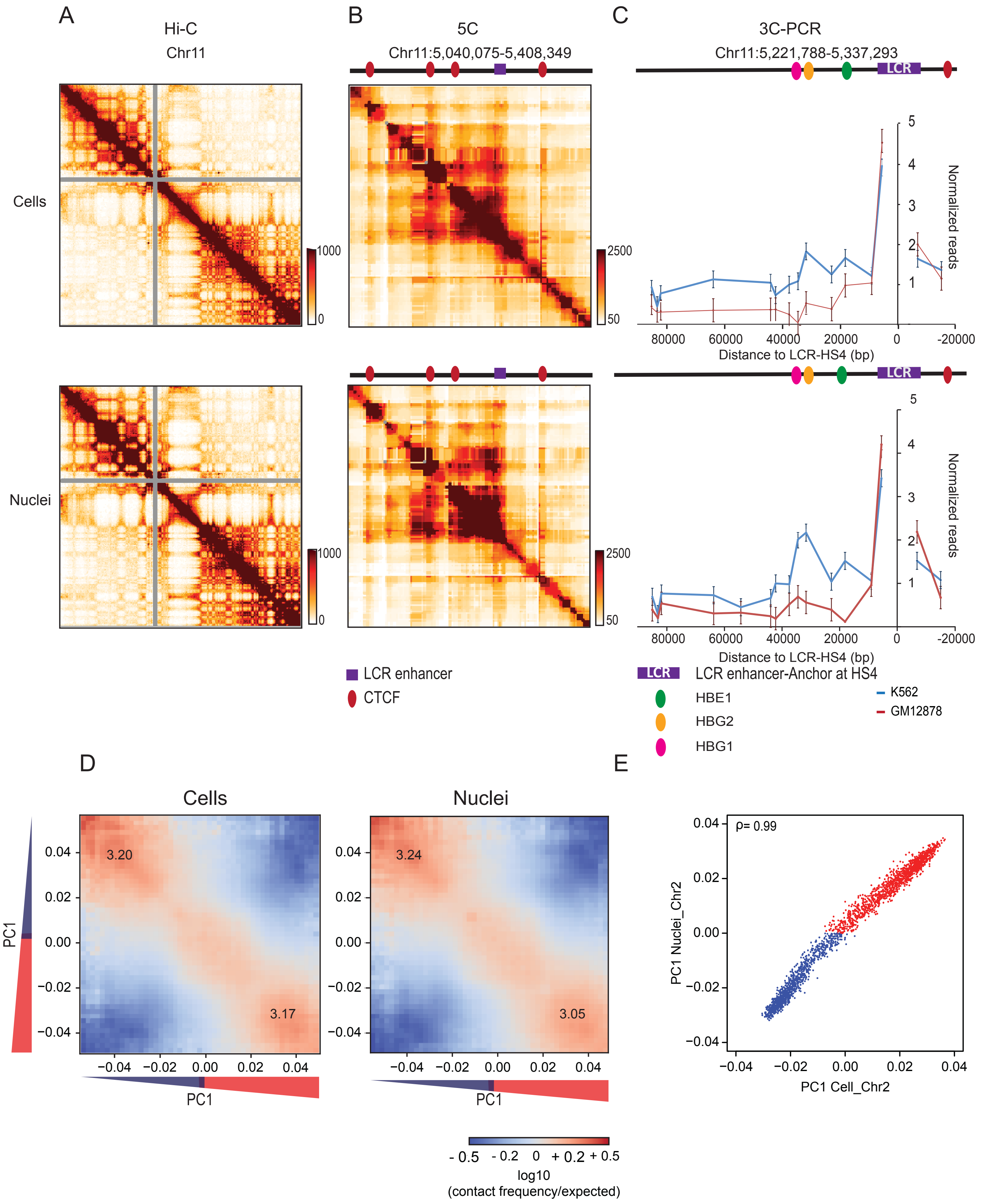

**Figure S1, related to Figures 1 and 2: Chromosome conformation in isolated nuclei**

**(A)** Hi-C 2.0 intra-chromosomal interaction maps for K562 cells display chromosomal compartments and TADs. Top: cells. Bottom: purified nuclei.

**(B)** 5C interaction map of 1 Mb region surrounding the beta-globin locus in K562 cells. Top: cells. Bottom: purified nuclei. CTCF-mediated interactions are preserved in purified nuclei. Red circles: positions of CTCF sites, purple square Beta-globin locus control region (LCR).

**(C)** 3C-PCR for a 44120 kb region surrounding the beta-globin LCR on chromosome 11, detects at high resolution the known looping interactions between the LCR and the expressed gamma-globin genes (HBE1, HBG2) in K562 cells. Looping interactions are not detected in GM12878 cells that do not express these genes. Top: cells. Bottom: purified nuclei.

**(D)** Compartmentalization saddle plots: average intra-chromosomal interaction frequencies between 100 kb bins, normalized by genomic distance. Bins are sorted by their PC1 value derived from Hi-C data obtained with K562 cells. In these plots preferential B-B interactions are in the upper left corner, and preferential A-A interactions are in the lower right corner. Numbers in the corners represent the strength of AA interactions as compared to AB interactions and BB interactions over BA interactions. Left: cells. Right: purified nuclei.

**(E)** Spearman correlation ($\rho$) of PC1 in cells vs PC1 in nuclei for chromosome 2 at 100kb resolution ($\rho$= 0.99).

Supplemental Figure 2

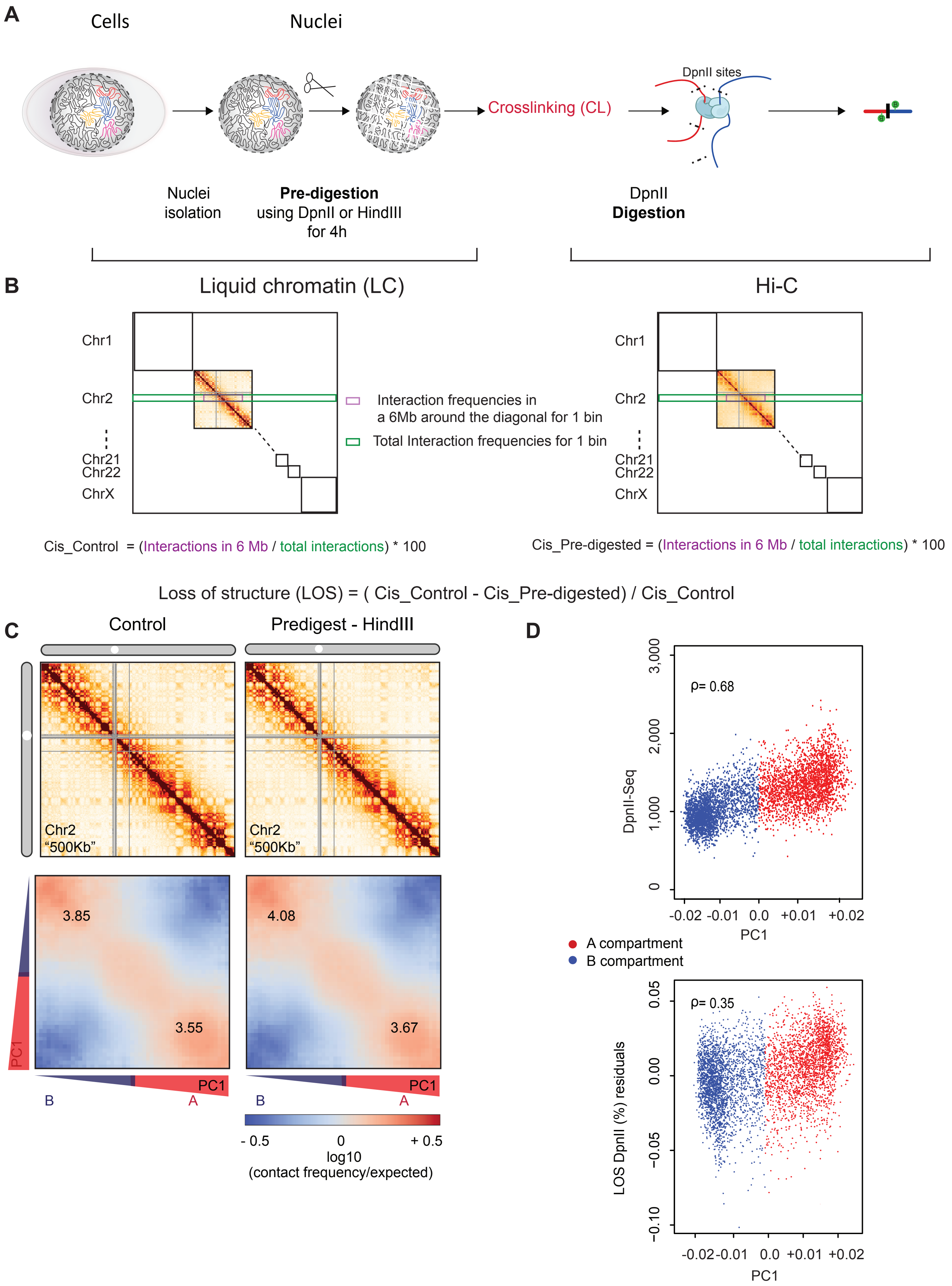

**Figure S2, related to Figure 3: Chromosome conformation dissolution upon chromatin fragmentation**

**(A)** Workflow for Liquid chromatin Hi-C.

**(B)** Illustration of loss of structure metric using a pre-digested sample and a control.

**(C)** Hi-C interaction maps and compartmentalization saddle plots for a second replicate of control nuclei (incubated for 4 hours in restriction buffer) and nuclei pre-digested with HindIII for 4 hours.

**(D)** Top: Spearman correlation of DpnII restriction digestion efficiency (DpnII-seq) and PC1 for chromosome 2 at 40 kb resolution. Bottom: Partial correlation of LOS (LOS residuals) with PC1 after controlling for restriction efficiency (DpnII-seq), for chromosome 2 at 40kb resolution. Spearman correlation is indicated.

Supplemental Figure 3

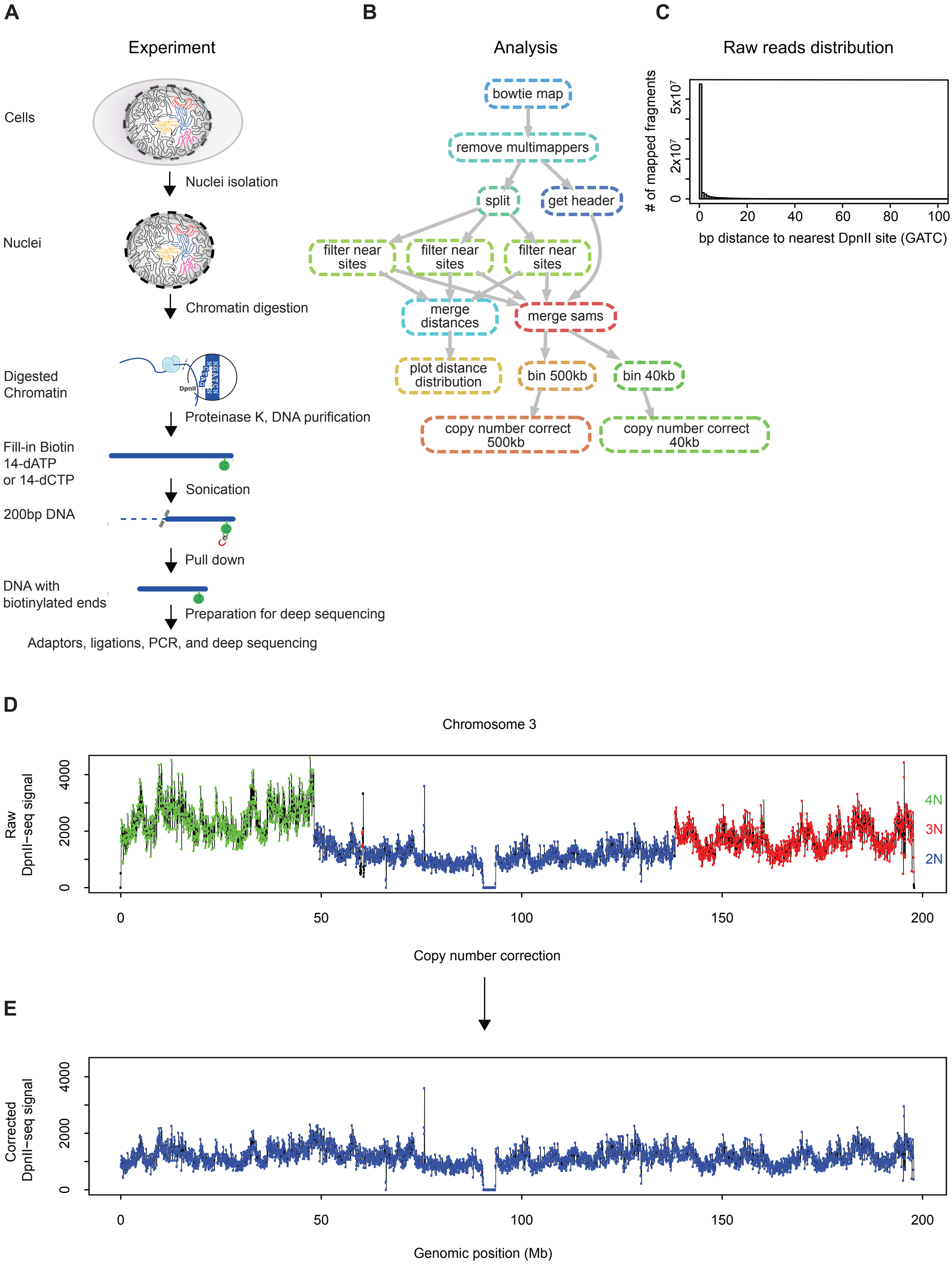

**Figure S3, related to Figure 3 and 4: Experimental protocol and computational workflow for DpnII-seq**

**(A)** Schematic of DpnII-seq experimental protocol for recovering DNA fragments digested by the restriction enzyme DpnII.

**(B)** Directed graph of DpnII-seq computational pipeline

**(C)** Histogram of distance to nearest DpnII recognition site for each recovered DpnII digested fragment.

**(D)** Raw DpnII-seq signal displaying multiple copy number states (2N, 3N, 4N) within chromosome 3 (data binned at 40 kb).

**(E)** Copy number corrected DpnII-seq signal displaying single copy number state (2N) across chromosome 3

Supplemental Figure 4

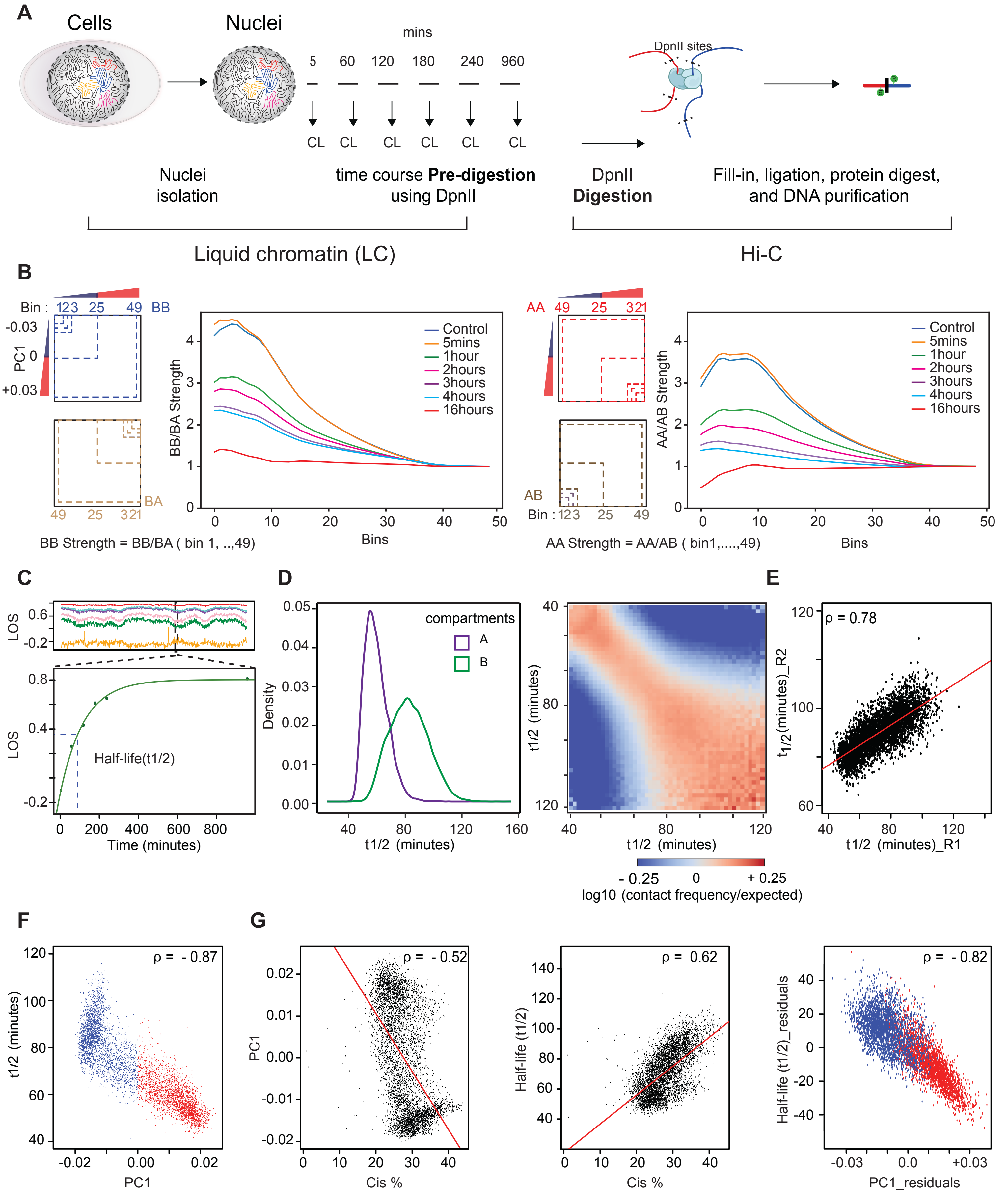

**Figure S4, related to Figure 4: Liquid chromatin-Hi-C protocol and quantification of loss of structure after chromatin pre-digestion**

**(A)** Workflow for Liquid chromatin Hi-C timecourse. CL = cross-linking step.

**(B)** Compartment strength derived from compartment saddle plots (See Methods). Left: Diagram depicting compartment strength calculation for B-B interactions. Plot to the right of diagram: B-B interaction strength as a function of bin number for all timepoints of the time course. Right: Diagram depicting compartment strength calculation for A-A interactions. Plot to the right of diagram: A-A interaction strength as a function of bin number for all time points of the time course.

**(C)** Top: LOS signal across a 40 Mb region on chromosome 2 calculated for indicated timepoints in the digestion timecourse. Line colors as in Figure 4E. Bottom: Exponential curve fit to LOS timepoints for a single 40kb bin. $t_{1/2}$ (dashed vertical blue line) representing time elapsed to reach half saturation of LOS signal.

**(D)** Left: Density distributions of $t_{1/2}$ for A and B compartments. Right: $t_{1/2}$ saddle plots: average intra-chromosomal interaction frequencies between 40 kb bins, normalized by genomic distance. Bins are sorted by their $t_{1/2}$ value derived from digestion timecourse. Bins with high $t_{1/2}$ preferentially interact (bottom right of heatmap) and bins with low $t_{1/2}$ preferentially interact (top left of heatmap).

**(E)** Scatterplot of $t_{1/2}$ vs $t_{1/2}$ for two timecourse replicates (R1 and R2) on chromosome 2. Regression line (red). Spearman correlation is indicated.

**(F)** Scatterplot of PC1 vs $t_{1/2}$ for chromosome 2. A compartment (red); B compartment (blue).

**(G)** Left: Scatterplot of percent interactions occurring *in cis* within a 6 Mb distance out of total genome wide interactions for each 40 kb bin in control Hi-C map (Cis %) vs PC1. Middle: Cis% vs $t_{1/2}$. Right: Scatterplot of partial correlation between PC1 and $t_{1/2}$ controlled by Cis %. A compartment (red); B compartment (blue). Solid red lines are regression lines. Spearman correlations are indicated.

Supplemental Figure 5

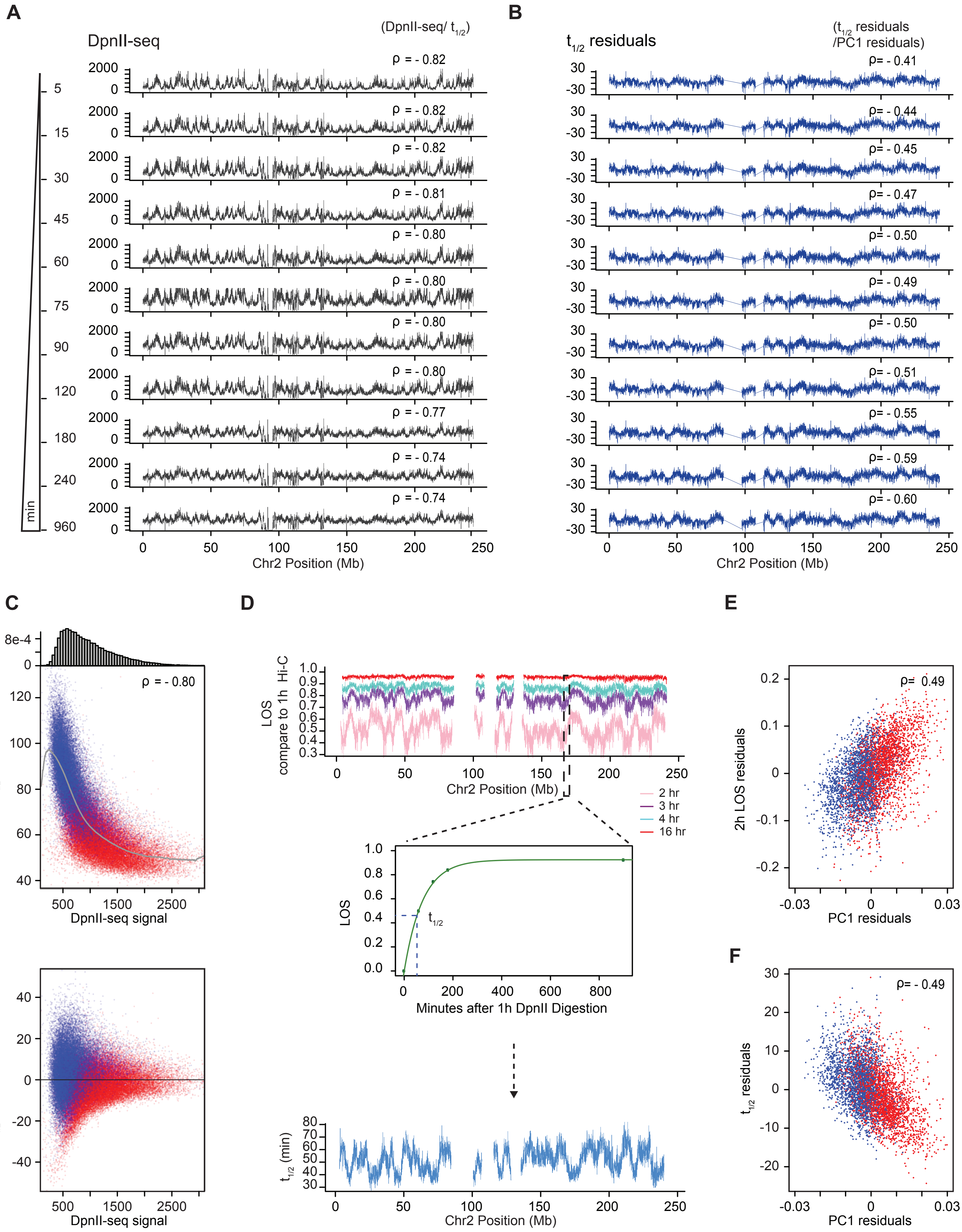

**Figure S5, related to Figure 4: Residual variation in half-life and LOS after correcting for DpnII cutting frequency is correlated with compartment state**

**(A)** DpnII-seq signals along chromosome 2 after indicated times of digestion. Spearman correlations between DpnII-seq and $t_{1/2}$ at each timepoint is indicated.

**(B)** $t_{1/2}$ residuals along chromosome 2 after correcting $t_{1/2}$ values by the correlation between $t_{1/2}$ and DpnII-seq signals shown on the left obtained after the indicated times of digestion. Spearman correlation between $t_{1/2}$ residuals and PC1 residuals are indicated.

**(C)** Top: Genome wide scatterplot of $t_{1/2}$ versus 1 hour DpnII-seq signal. Gray line: moving average. Bar plot above shows the number of loci displaying various levels of DpnII-mediated cuts. Bottom: residuals of $t_{1/2}$ calculated by subtracting $t_{1/2}$ from the corresponding average $t_{1/2}$ (gray line in top plot) plotted vs. number of DpnII cuts. Red dots: loci in the A compartment; Blue dots: loci in the B compartment. The majority of loci have 500-1100 cuts. When comparing loci with similar number of DpnII cut we observe that loci in the A compartment have shorter $t_{1/2}$ values as compared to loci in the B compartment.

**(D)** Top: LOS along chromosome 2 at the indicated timepoints of digestion and calculated by comparison to Hi-C data obtained after 1 hour of digestion. Middle: calculation of $t_{1/2}$ from LOS at different timepoints. Bottom: $t_{1/2}$ along chromosome 2. This $t_{1/2}$ is calculated using the Hi-C data obtained after 1 hour of pre-digestion as starting point.

**(E)** Partial correlation between LOS and PC1 after correcting for their correlations with DpnII-seq. LOS (at 2 hours) is calculated as in panel C using the Hi-C data obtained after 1 hour of pre-digestion as starting point

**(F)** Partial correlation between $t_{1/2}$ and PC1 after correcting for their correlations with DpnII seq. $t_{1/2}$ is calculated as in panel D using the Hi-C data obtained after 1 hour of pre-digestion as starting point. Spearman correlations are indicated.

### Supplemental Figure 6

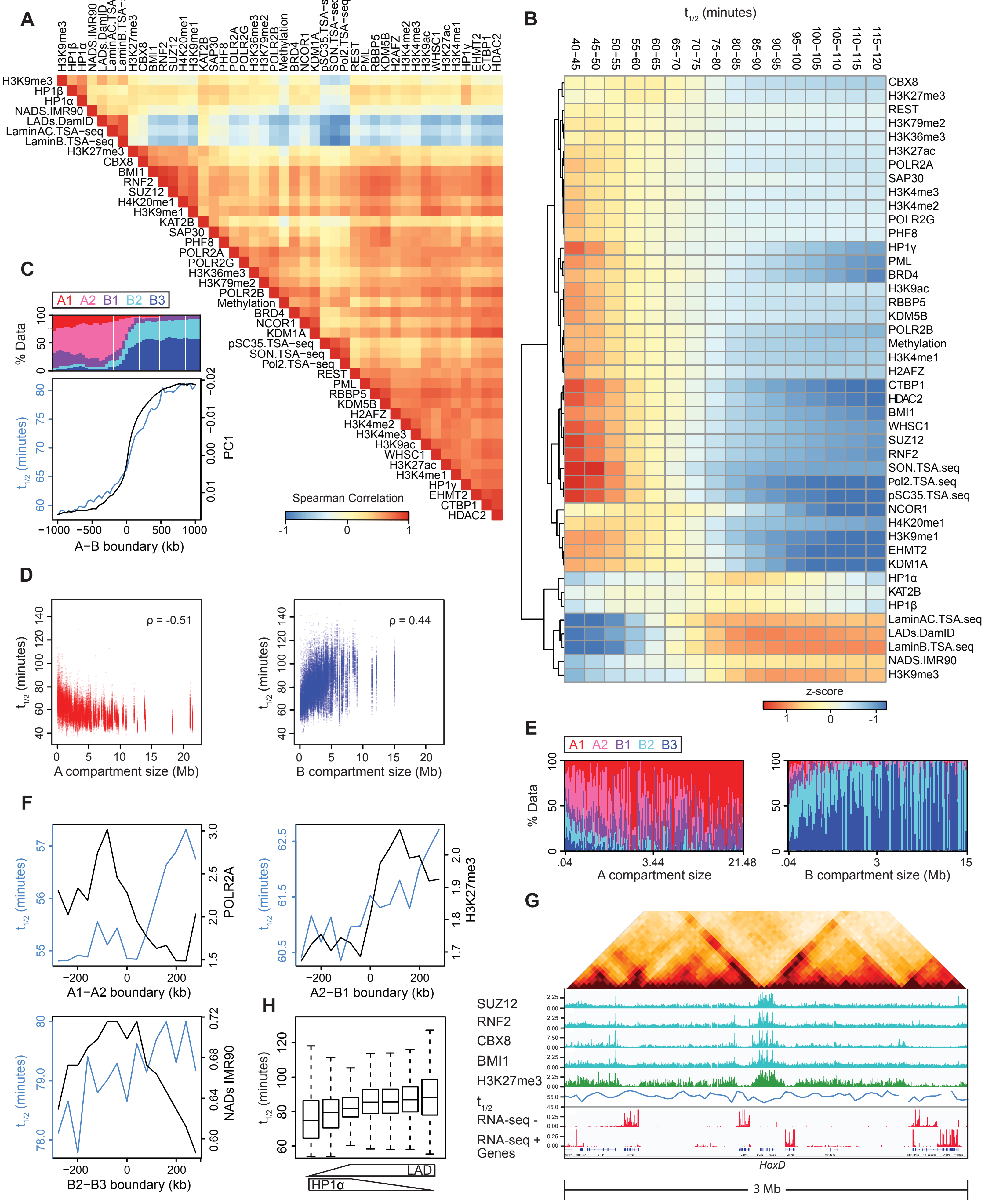

**Figure S6, related to Figure 4 and 5: Associations between sub-nuclear structures, compartment characteristics and chromatin interaction stability**

**(A)** Spearman correlation matrix between signals for various chromatin state markers of various sub-nuclear structures, chromatin remodellers and histone modifications with row order determined by hierarchical clustering.

**(B)** The genome was split into 16 bins, where each bin corresponds to sets of loci that share the same $t_{1/2}$ interval. For each $t_{1/2}$ interval the mean z-score signal enrichment for various markers of sub-nuclear structures, chromatin remodellers and histone modifications was calculated and shown as a heatmap. Row order determined by hierarchical clustering.

**(C)** Top: Stacked barplot displaying percentages for each of five sub-compartments within 1 Mb of A to B boundary switches with flanking A/B compartments larger than 1 Mb. Bottom: Transitions of mean $t_{1/2}$ (blue) and mean PC1 (black) at same A to B boundary switches.

**(D)** Left: Scatterplot of $t_{1/2}$ vs size of A compartment containing $t_{1/2}$ bin. Right: $t_{1/2}$ vs size of B compartment containing $t_{1/2}$ bin.

**(E)** Left: Stacked barplot displaying percentages for each of five sub-compartments across and within A compartments of a specific size. Right: Stacked barplot displaying percentages for each of five sub-compartments across and within B compartments of a specific size.

**(F)** Transitions of mean $t_{1/2}$ (blue) and mean chromatin feature (black) across various sub-compartment boundary switches.

**(G)** 3 Mb region surrounding *HoxD* locus. Top: Hi-C contact map for K562 control nuclei showing the position of the *HoxD* locus. Tracks: ChIP-seq tracks for polycomb subunits (cyan) and the polycomb associated histone modification H3K27me3 (green). $t_{1/2}$ (blue). Minus strand and plus-strand signal of total RNA-seq (red). Refseq Genes (blue/black). The polycomb-bound domain displays shorter half-life compared to expressed genes in flanking regions.

**(H)** Boxplots of $t_{1/2}$ stratified by bins enriched in HP1α and depleted in LADs (left) and bins depleted in HP1α and enriched in LADs (right).

**Supplemental Table S1, related to Supplemental Figure S1.** Sequences of 3C primers

**Supplemental Table S2, related to Supplemental Figure S1.** Sequences of 5C primers

**Supplemental Table S3, related to Figure 5 and Supplemental Figure S5.** Public datasets used in this paper.

**Supplemental Movie 1, related to Figure 1.** Force extension experiment for a control nucleus.

**Supplemental Movie 2, related to Figure 2.** Force extension experiment for a nucleus digested with DpnII.
