## Supplementary figures and images for "Compartment-dependent chromatin interaction dynamics revealed by liquid chromatin Hi-C"

### Supplemental Movies

#### Slide 1
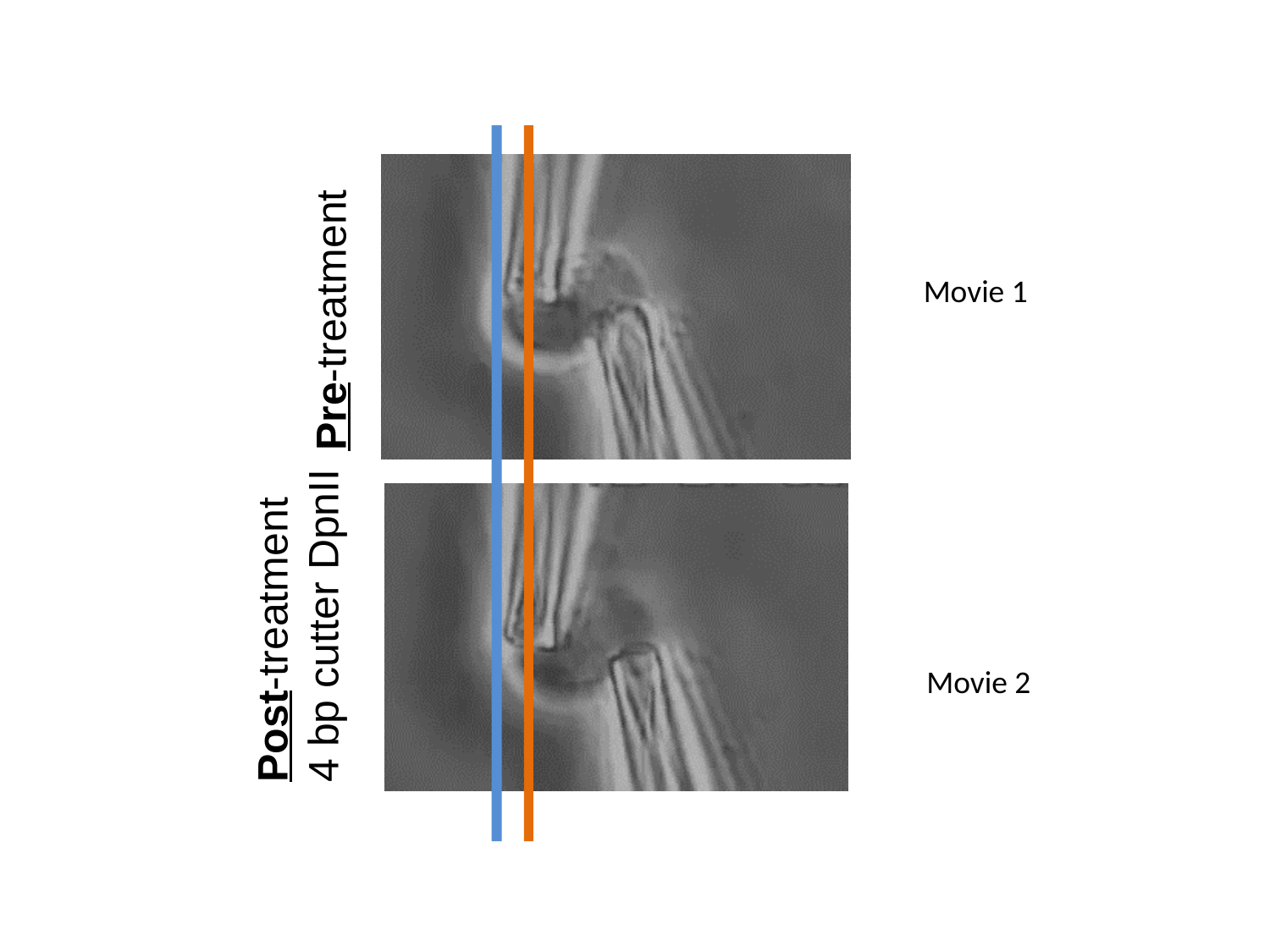

Movie 1
Pre-treatment
Post-treatment
4 bp cutter DpnII
Movie 2
